# Computational Identification of Migrating T cells in Spatial Transcriptomics Data

**DOI:** 10.1101/2024.10.23.619870

**Authors:** Lin Zhong, Bo Li, Siyuan Zhang, Qiwei Li, Guanghua Xiao

## Abstract

T cells are the central players in antitumor immunity, and effective tumor killing depends on their ability to infiltrate into the tumor microenvironment (TME) while maintaining normal cytotoxicity. However, late-stage tumors develop immunosuppressive mechanisms that impede T cell movement and induce exhaustion. Investigating T cell migration in human tumors in vivo could provide novel insights into tumor immune escape, although it remains a challenging task. In this study, we developed ReMiTT, a computational method that leverages spatial transcriptomics data to track T cell migration patterns within tumor tissue. Applying ReMiTT to multiple tumor samples, we identified potential migration trails. On these trails, chemokines that promote T-cell trafficking display an increasing trend. Additionally, we identified key genes and pathways enriched on these migration trails, including those involved in cytoskeleton rearrangement, leukocyte chemotaxis, cell adhesion, leukocyte migration, and extracellular matrix (ECM) remodeling. Furthermore, we characterized the phenotypes of T cells along these trails, showing that the migrating T cells are highly proliferative. Our findings introduce a novel approach for studying T cell migration and interactions within the tumor microenvironment (TME), offering valuable insights into tumor-immune dynamics.

## Introduction

T cell cytotoxicity is essential for the anti-tumor adaptive immune response(*1, 2*). After primed by tumor antigens in the draining lymph spots(*3*), effector T cells travel through the lymphatic vessels to enter the tumor microenvironment (TME)(*4*). This process is mediated by the T cells expressing necessary cell adhesion molecules and chemokine receptors for migration and infiltration in the tumor(*5, 6*). Efficient tumor killing hinges on 1) the ability of T cells to infiltrate into the tumor core regions(*7*) and 2) T cells maintaining a high level of functionality and cytotoxicity. In late-stage tumors, however, either or both processes were disabled, creating a hostile environment to prevent T cell infiltration(*8*), or rapidly inducing T cell exhaustion during their migration inside the tumor region(*9*).

Clinical efforts have been prioritized to lift T cell exhaustion by immune checkpoint blockade(*10, 11*), or to break down the tumor barrier to turn immune ‘cold’ tumors ‘hot’(*12*). In addition, reduced motility has been observed among exhausted T cells(*13*).Therefore, investigation on how T cell migrate inside the tumor region and how they become dysfunctional along the way may provide novel therapeutic insights. To date, *in vitro* experiments have been performed to track T cell movement on 2-dimensional (2D) or 3D culture conditions, which improved the quantitative understanding of T cell migration(*14–17*). Non-invasive *in vivo* tracing of cell movement requires the use of radioactive or fluorescent dyes which can be monitored with intravital imaging. This has been done using transgenic mouse models with high single-cell resolution (*18, 19*). However, this approach is usually technically demanding and cannot reach the resolution and scale to study T cell migration in the TME. Consequently, there is still a lack of investigations on T cell migration in the TME.

The recent development of the cutting-edge spatial transcriptomics technologies, such as 10xVisium, MERFISH, Slide-Seq, NanoString CosMx, etc, (*20–23*) provided a new opportunity to address this problem. Spatial transcriptomics data maps the location of gene expression directly within a tissue’s spatial architecture and allows researchers to study gene activity within the context of tissue structure. In this work, we developed a novel computational method, ReMiTT (Reconstruction of Migration Trails for T cells), to identify T cell migration trails using spatial transcriptomics data. Previous in vitro studies demonstrate that when navigating through a 3-D collagen matrix, T cells prefer to migrate through less dense or channel-like regions, often following each other along the same trajectory (15). We hypothesize that specific areas with remodeled tumor extracellular matrix (ECM) serve as physical highways facilitating T cell migration. Considering that T cells inevitably become exhausted in solid tumors, an increasing trend of exhaustion levels may be observed along some of these migration trails, where streams of T cells follow each other to enter and traverse these regions. ReMiTT is designed to locate these trafficking T cells by searching for closely located T cell loci or spots that exhibit an increasing trend of exhaustion. This might be a snapshot of a stream of migrating T cells on the “physical highway”. We implemented this method to study human tumor samples and identified multiple T cell migration trails in a tissue slide. The biological relevance of the trails was validated using T cell chemotaxis markers and T cell receptor genes. Systematic investigations of the genes overexpressed on the migration trails show enrichment of gene pathways related to T cell migration, chemokine production, cytoskeleton rearrangement, and ECM remodeling on these trails. Furthermore, we identified evidence that immunosuppressive cell types, such as Treg have elevated levels on migration trails and density of tumor-associated macrophages increases along the trails. Our study might provide a novel approach to unbiasedly explore T cell migration patterns and related genes in human cancer samples.

## Results

### Characterization of T cell migration trails

We implemented ReMiTT to analyze tumor samples profiled using the 10x Visium spatial transcriptomics platform. This approach is based on a simple observation that migrating T cells in the tumor microenvironment will gradually lose their ability to kill the cancer cells and enter an exhaustion state (*24*). Hence, we hypothesized that the continued movement of T cells can be captured by ‘stitching’ neighboring spots on a 2-D slide with matched T cell phenotypes, which leads to the development of ReMiTT. In brief, we first identify spots that are most likely to contain newly entry T cells from the draining lymph spots as the ‘start point’. ReMiTT then searches for 1D trails on a 2D slide such that T cells on these trails display an increasing trend of a pre-defined score of T cell exhaustion, using either a minimum spanning path (MSP) searching method or a 3-D line-based approach (**Figure 1**).

**Fig. 1.**
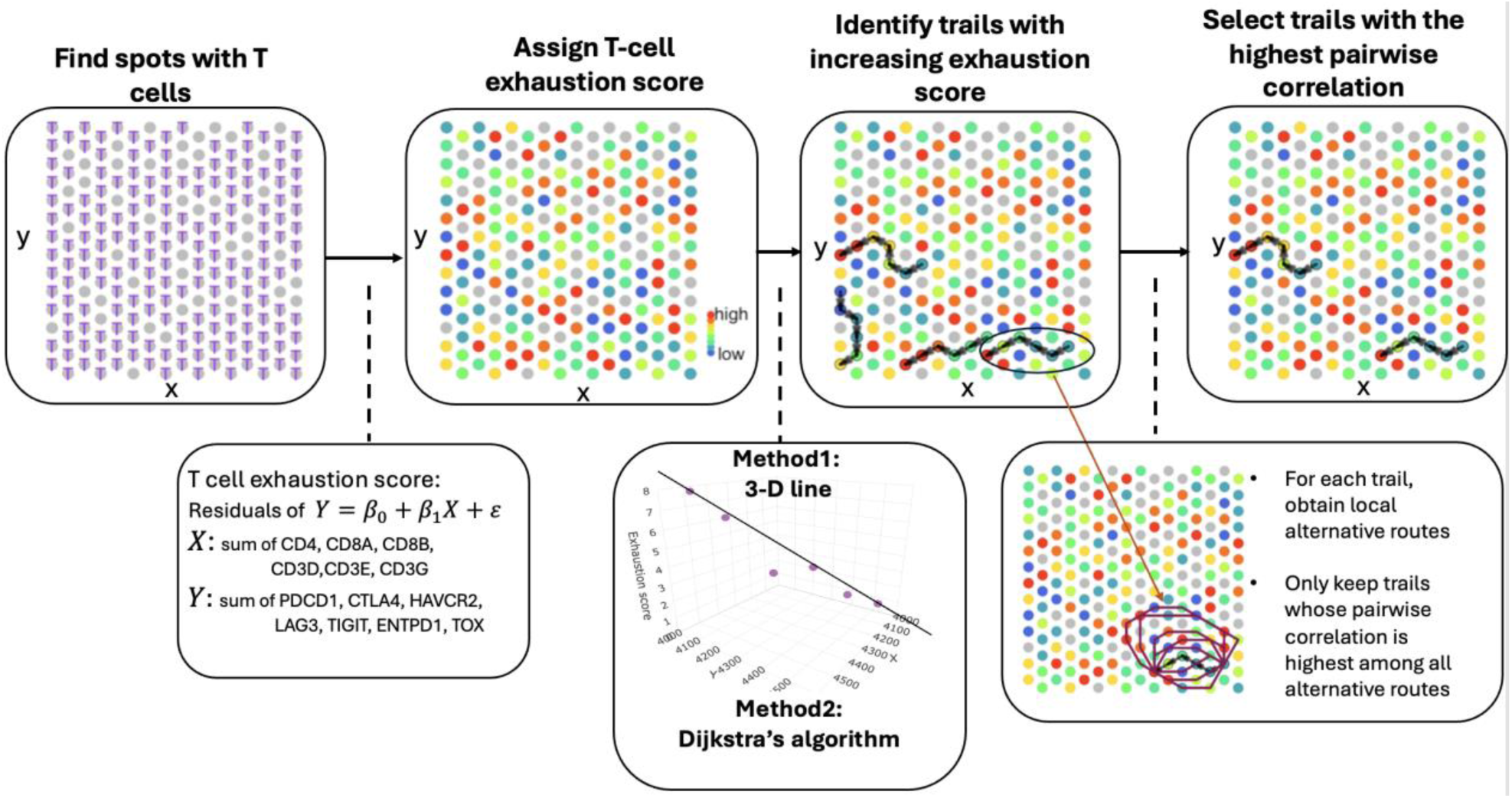
Schematic illustration of the T cell migration trail identification method. We first identified spots with expression of T cell surface markers (CD4, CD8 and CD3) to select those with T cell infiltration. A T cell exhaustion score was then assigned for each T cell spot using putative exhaustion markers. Candidate migration trails were identified by searching for increasing exhaustion scores among adjacent spots. To filter for natural migration trails, we only keep the ones whose mean pairwise correlation among genes associated with T cell stage is higher compared to all its alternative routes.

We applied ReMiTT to a human lung tumor sample and identified 19 T-cell migration trails (**Figure 2A**). The trail lengths ranged from 6 to 9 spots, with an average span of 7 spots (∼0.7 mm). Notably, some trails were observed to originate near blood vessels (**Figure 2B-C**), while others predominantly traversed along loosened tissue regions (**Figure 2D**). These findings align with our hypothesis that T cells may preferentially navigate through less dense areas within the tumor microenvironment (TME) to avoid physical barriers such as abnormally dense collagen fibers in tumor that trap them. Of the 19 migration trails, three extended deep into the tumor core, with a median length of 0.7 mm. This suggests that T cells might rapidly become exhausted after relatively short journeys into the immunosuppressive TME.

**Fig. 2.**
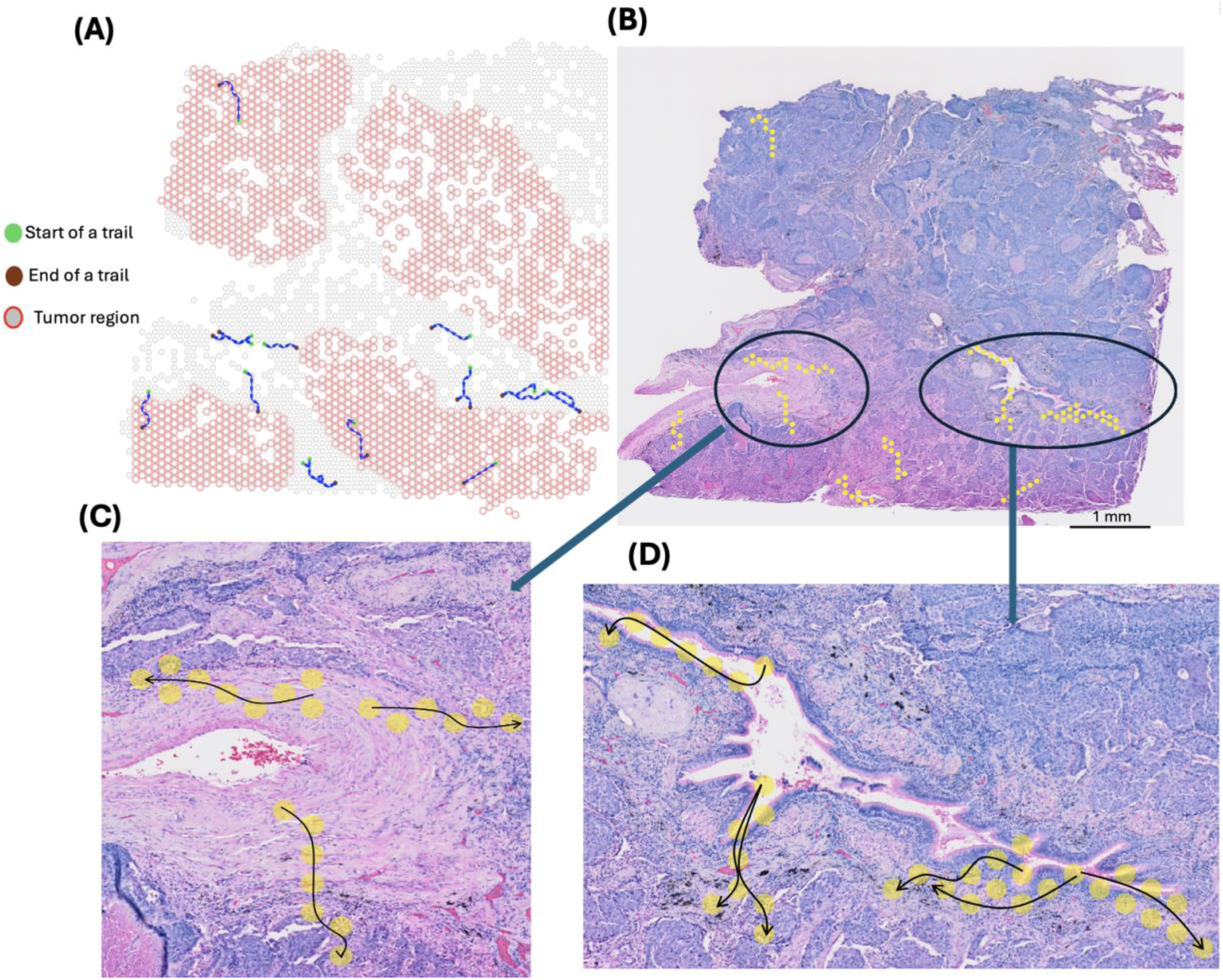
Migration trails identified in a human lung cancer sample and their projections on the pathology slide. **(A)**Algorithm-identified migration trails labeled as splines on the spatial coordinates, with the starting location marked as green. Grey spots are spots with T cell infiltration and red circles indicating spots with high levels of expression of ovarian cancer markers. **(B)** Same trails overlaid on the pathology (H&E) slide of the tumor. **(C)** Examples of trails whose origin is close to vessels **(D)** Examples of trails travel in less dense TME.

### Elevated T cell chemotaxis along the migration trails

As experimental tracing of T cell migration *in vivo* is challenging, we sought to validate the trails through biological findings. Specifically, T cell migration is regulated by chemokine/receptors in the tumor microenvironment (*6*). Several key chemokine receptors, including, CXCR3/4/6, etc are reported to direct T cell migration in the tumor (*6, 25*). In the meanwhile, it is known that T cells respond to a positive gradient of the chemoattractants that guides the movement to the loci of interest (*26, 27*). We investigated the chemokines associated with putative T cell migration regulators, including CXCL9/10/11/16 and CCL4/5 along the migrating trails. These ligands have been reported to promote T cell infiltration into tumor(*25*). We observed a significantly increasing trend for all the CXC motif chemokines, including CXCL9/10/11 (ligand for CXCR3) and CXCL16 (ligand for CXCR6), whereas the trend for CCL4/5 was not significant in this lung cancer sample (**Figure 3**). CXCR3 is one of the most well-studied receptor for T cell migration (*28, 29*), and has been implicated in antigen-specific T cell responses (*30*). No trend was observed for the chemokine receptors on the trails, possibly because the expression of these receptors is sufficient to induce T cell migration (*6*). We made similar observations in a melanoma sample and an ovarian tumor sample (Supplementary Figure 1). These results supported our findings of the T cell migration trails using ReMiTT.

**Figure 3.**
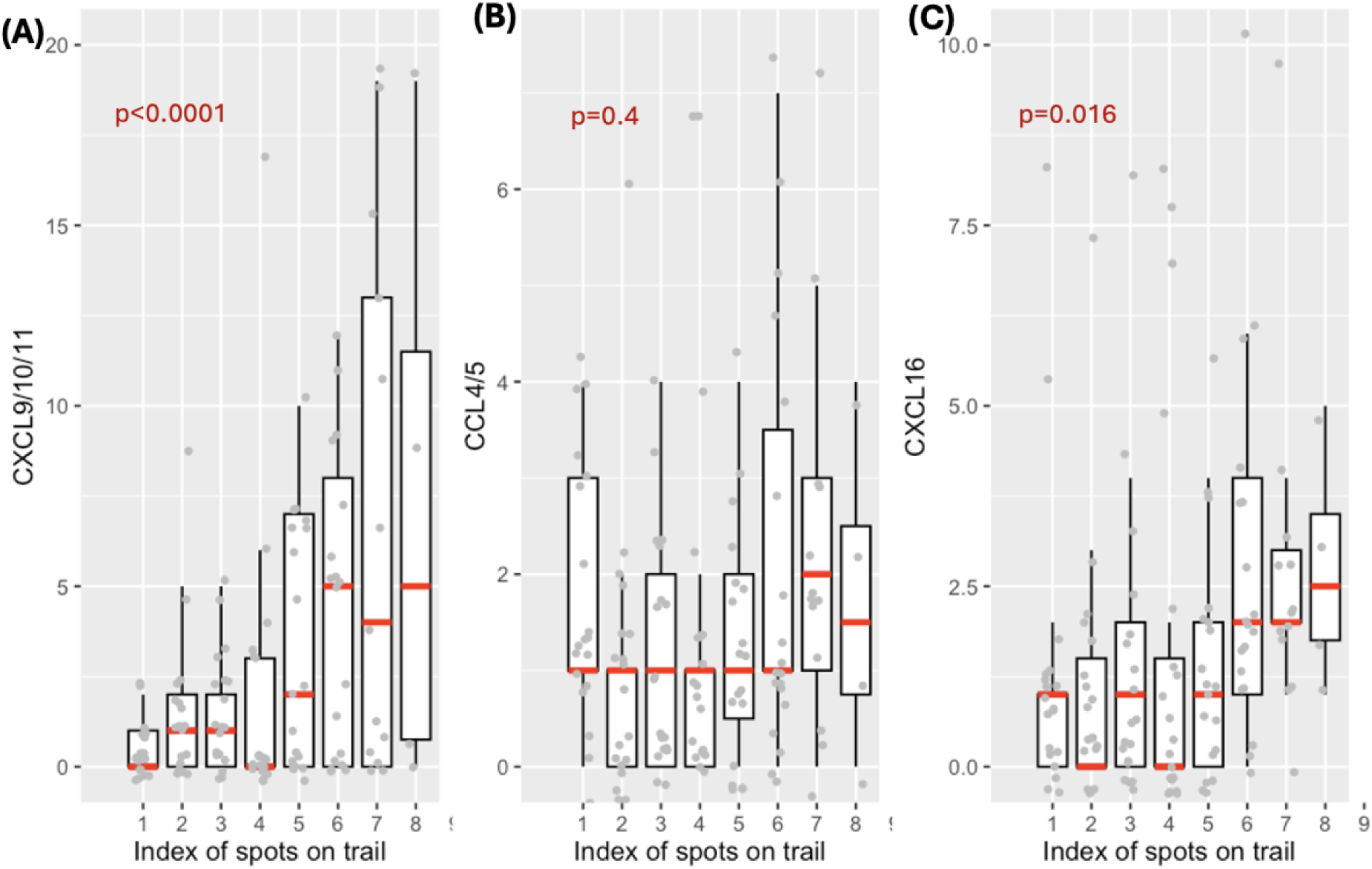
Trend of chemokines that promote T cell migration along the algorithm-identified migration trails, and phenotype of these trails identified in the lung cancer sample. **(A)**Total expression of CXCL9, CXCL10, and CXCL11 along the algorithm identified migration trails with the median of each index labeled with a red line segment. Mixed-effect linear regression adjusting for UMI of each spot is performed to test the increasing trend of ligand expression along the trails, and 2-sided p value is reported in the panel. **(B)** Total expression of CCL4 and CCL5 along the algorithm identified migration trails, with the median of each index labeled with a red line segment. **(C)** Expression of CXCL16 along the algorithm identified migration trails,with the median of each index labeled with a red line segment.

### Shared TCR genes between spots on trail

Next, we sought to provide another validation using a statistical approach. T cell antigen priming and proliferation occur in the draining lymph spots(*31*), prior to the migration to the tumor site. Hence, it is expected that T cells of the same clone are more likely to share a migration trajectory. Here, as paired TCR hypervariable regions were not sequenced, we used the TCR α variable (TRAV) and joining (TRAJ) genes and the TCR β variable (TRBV) genes as surrogates to trace clonal sharing. The Visium platform for spatial genomic profiling comes in two versions—v1 (original) and v2 (enhanced), and the TCR variable genes are only captured in v1, but not v2, which is used for our lung sample. Thus, we utilized an ovarian cancer sample from v1 for this analysis. To test whether spots on migration trails shared more TCR variable genes compared to spots not on trails, we first obtained one matching trail for each of the algorithm-identified migration trails. We call it one control set. We then obtained 5,000 such matched control sets. An example of one control set is displayed in Supplementary Figure 2. The systematic analysis in this study is then based on comparing algorithm-identified migration trails with their matched control trail sets, rather than simply comparing spots on and off the migration trails. This approach accounts for the spatial distribution of gene expression, as requiring a ‘trail’ to only consist of adjacent spots influences the expected gene expression pattern.

We calculated the mean number of shared TCR V/J genes among consecutive pairs of spots along each of the algorithm-identified trails. We then calculated the same statistics for each of the 5,000 control sets and got the empirical distribution of the median of the mean shared TCR V/J genes of each trail for each control set. (Supplementary Figure 3). This works as a surrogate for the shared TCR of each trail. The median number of the shared TCR V/J genes among the 19 migration trails is significantly higher (median=13.6) compared to the median in the empirical distribution of these statistics in control sets (median=11.2), with the 2-sided empirical p-value being 0.005. As different immune cells are reported to migrate together(*6*), we also investigated the clonal sharing of B cells, under the premise that the physical passages for T cell migration might also be utilized by B cells. Consistently, we observed that the median number of shared BCR variable genes among the 19 migration trails is significantly higher (median=2.4) compared to the median in the empirical distribution (median=1.9), with the 2-sided empirical p-value is 0 (Supplementary Figure 3). While these results may not provide direct validation, they supported our conclusion that the predicted trails are statistically enriched for immune cell migration paths.

### Gene expression signatures along the migration trails

We next investigated the gene expression signatures of cells on the migrating trails. We performed differential gene expression analysis by comparing the mean expression levels of the 3000 variable genes in this lung cancer sample on the migration trails to the empirical distribution of gene expression generated with the 5,000 matched control trail sets. We identified 394 significantly upregulated genes (empirical FDR<0.05 and fold change >1.2, **Figure 4A**, **Supplementary Table 1**). Consistently, extracellular enzymes that break down collagen network, including MMP1 and MMP2 (*32*), were significantly upregulated on the trails. In addition, genes encoding cell adhesion molecules that shown to promote cell migration and matrix remodeling were also upregulated on the migration trails, such as ITGA8/11, TJP3, etc (*33–36*). Furthermore, we observed that LGALS3, which encodes Galectin-3, is significantly upregulated (adjusted p value=0.013). It is a ligand of LAG3 that induces reduced T cell cytotoxicity and exhaustion (*37*). This gene independently emerged as a top hit in our analysis, where LAG3 level has been controlled. This result indicated that Lag3/Galectin-3 axis might be responsible for inducing T cell exhaustion along the trails. To understand the gene pathways related to T cell migration, we performed gene-set enrichment analysis (GSEA) (*38*). As expected, the top hits contained many key components of cell migration (**Figure 4B-C**). These pathways included the ones involved in the regulation of immune cell chemotaxis, cytoskeleton rearrangement, cell adhesion, and extracellular matrix binding.

**Fig. 4.**
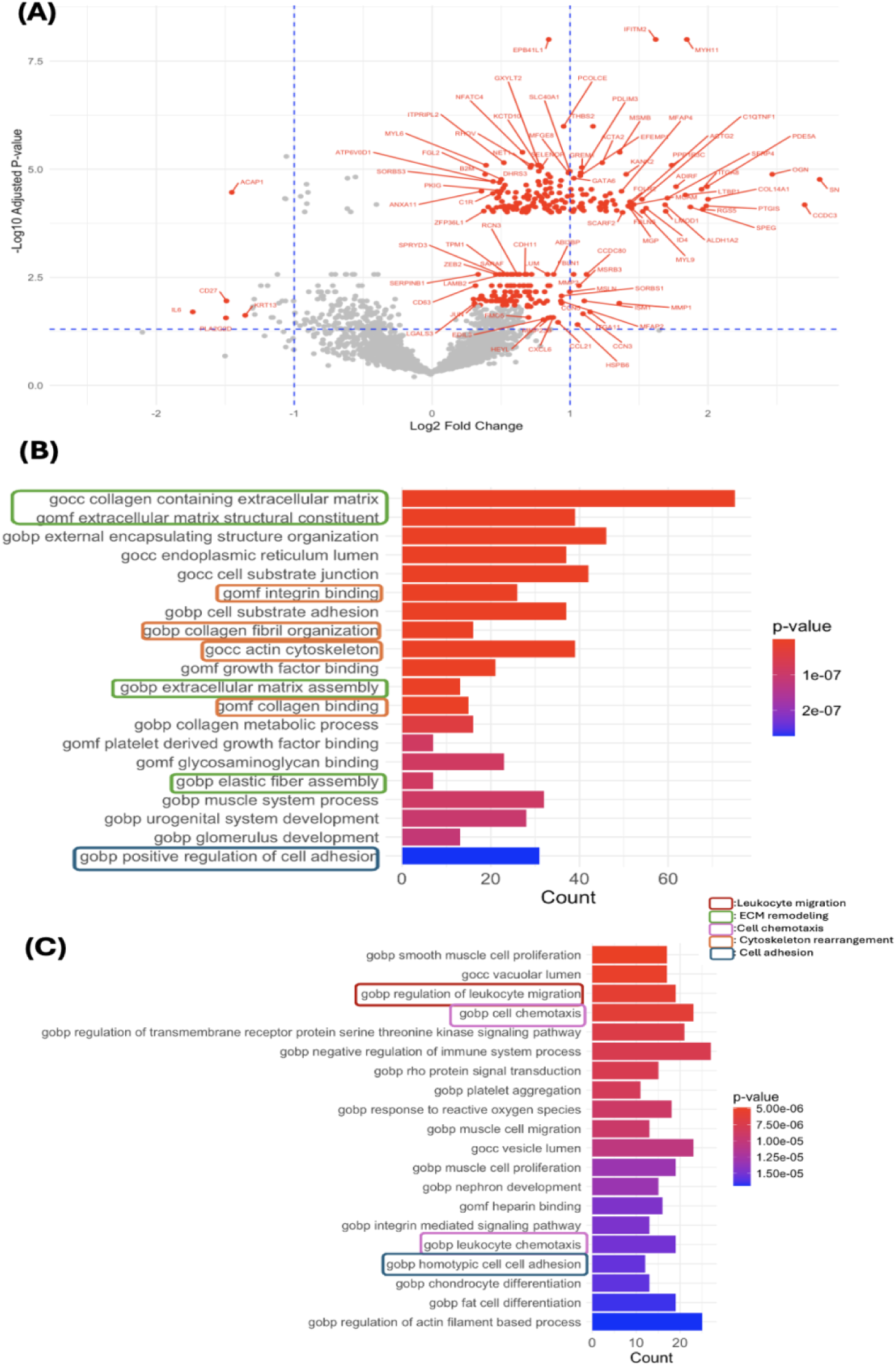
Differential gene expression analysis for T cell migration trails on the lung cancer sample. **(A)** Volcano plot of gene (3000 variable genes) fold change (FC) and adjusted empirical p values between those on migration trails vs. 10000 control sets. Genes with FDR<0.05 and the absolute values of FC ≥1.2 were labeled on the plot. **(B)** Barplot showing the top 20 enriched Gene Ontology biological process (GOBP) gene pathways computed from the 394 significantly upregulated genes on migration trails by GSEA. Colors of the textbox borders indicated the type of biological processes that may involved in T cell migration. Colors of the bars indicate the adjusted p values. Statistical significance was evaluated using Fisher’s exact test, with FDR corrected by the Benjamini-Hochberg approach. **(C)** Barplot showing the top 50-70 enriched Gene Ontology biological process (GOBP) gene pathways computed from the 394 significantly upregulated genes on migration trails by GSEA.

To validate the above observations, we analyzed another ovarian tumor sample generated with the 10x Visium spatial transcriptomics platform (Method). Implementing the same trail-identification method, we found 29 migration trails and most of them in the tumor region (**Supplementary Figure 4)**. The length of these trails varied between 6 to 8 spots with the average being 6.6 spots, similar to the first sample. For this sample, we also obtained 5,000 control trail sets, each containing 29 matched trails corresponding to the 29 algorithm-identified migration trails. We analyzed the upregulated genes on the migration trails and identified a total of 191 genes that are overexpressed along these trails compared with the control trails (**Supplementary Figure 5A**). Importantly, we reproduced the observation that Galectin-3 is upregulated on the migration trails (adjusted p value=0.010). We also observed similar GSEA results with pathways enriched for immune cell migration (**Supplementary Figure 5B-C**).

### Phenotype of on-trail migrating T cells

We next sought to explore the phenotypes of the T cells on the migrating trails using single-cell RNA-seq data profiled from human T cells. Specifically, we obtained the data from a previous study of 12,346 T cells from 14 non-small cell lung cancer patients(*39*). We used this dataset due to its high data quality and well-accepted T cell subset annotations. We performed unbiased clustering based on the top 30 PCs on these T cells and identified 13 clusters (**Figure 5A**). We compared genes upregulated on T cell migration trails to investigate which cluster showed similar gene expression signatures. We observed that cluster 9 and cluster 3 were top enriched (**Figure 5B**). This analysis was repeated using other tumor samples and we observed the same results (**Supplementary Figure 6**), suggesting that cluster 9 and 3 present the phenotype of the moving T cells on the migration trails.

**Fig. 5.**
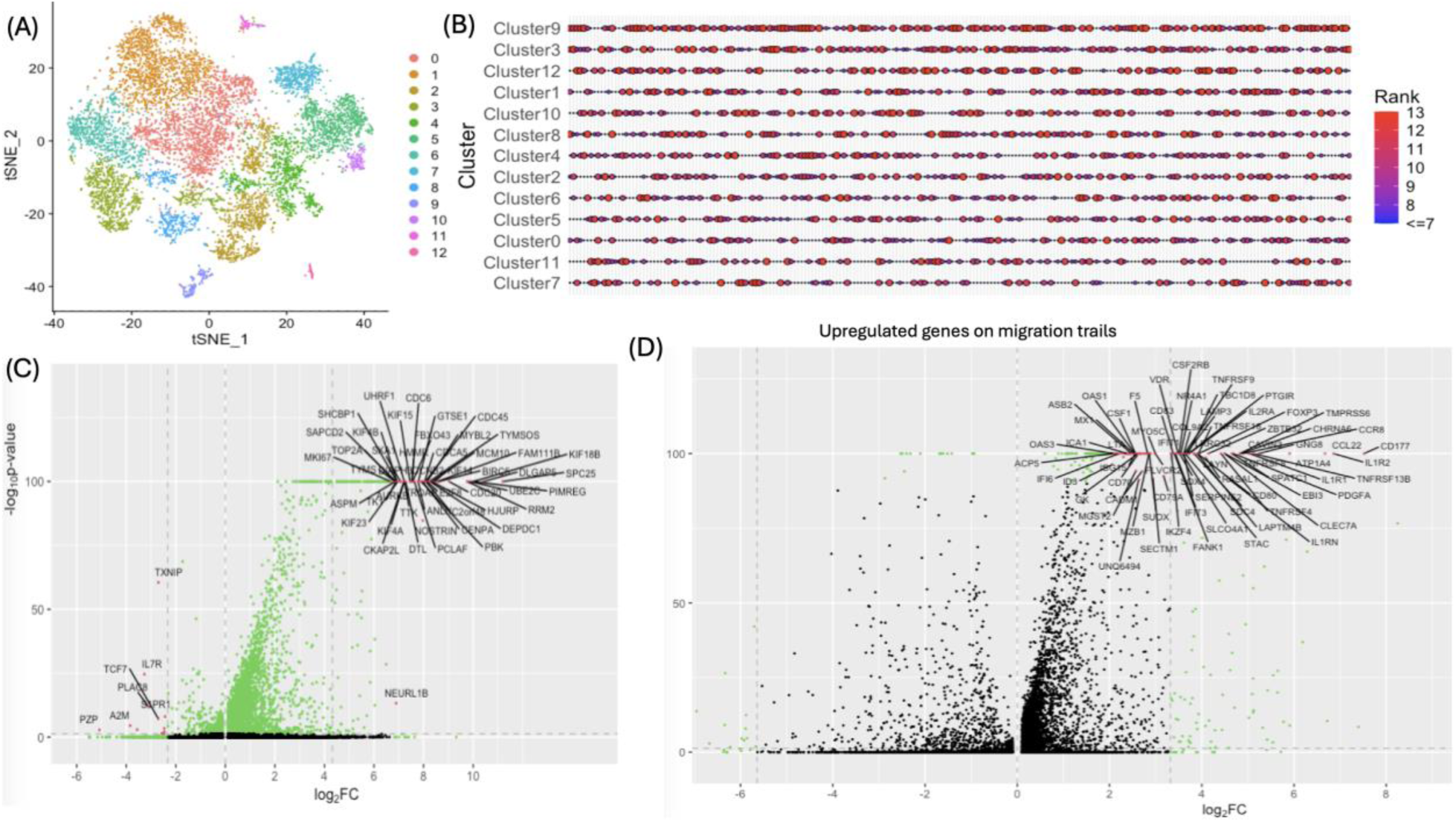
Phenotypic investigation of T cells on the migration trails. **(A)** t-SNE plot for the 11,236 T cells from deep single-cell RNA-seq data of 14 non-small lung cancer patients, with the nearest-neighboring clusters color-labeled. **(B)** Bubble plot showing the enrichment of overexpressed genes on T cell migration trails in each of the T cell clusters in (A). The large bubble sizes and red colors indicate higher ranks of mean expression of each of the upregulated genes on migration trails among all the 13 clusters of T cells. Clusters (y-axis) were ordered by their median ranks of the genes. **(C)** Volcano plots of gene fold change and adjusted p values for Cluster 9 compared with other clusters. Statistical significance was evaluated using the Wilcoxon rank sum test with FDR corrected using the Benjamini-Hochberg procedure. **(D)** Volcano plots of gene fold change and adjusted p values for Cluster 3 compared with other clusters. Statistical significance was evaluated using the Wilcoxon rank sum test with FDR corrected using the Benjamini-Hochberg procedure.

T cells of cluster 9 expressed high level of MKI67 and several members of the kinesin family (**Figure 5C**). These T cells are likely proliferating T cells with massive cytoskeleton remodeling and intracellular protein trafficking. We also identified cytotoxic markers in the top-upregulated genes in this cluster, suggesting that it might contain cytotoxic CD8 T cells. Cluster 3 expressed high levels of Treg markers FOXP3 and IL2RA (**Figure 5D**). These observations might be associated the a previously reported migrating Treg population (*40*).

This result also indicated that on the migration trails there may be a mixture of immune cell population. Hence, we also investigated tumor associated macrophages, which are also known regulators of T cell anti-tumor responses (*41*). Consistently, we observed an increasing trend of macrophage markers along the migration trails in melanoma and lung cancer (**Supplementary Figure 7**). Together, our results might suggest a co-migration of multiple immune cell types with T cells in the tumor microenvironment.

## Discussion

Tumor infiltrating lymphocytes (TILs) are the functional carriers of antitumor immunity. As T cells need to make physical contact with their target cells to perform the cytotoxic function, their successful infiltration into the tumor core is the basis of an effective immunotherapy of adoptive T cells(*42–46*). However, how T cells migrate within TME and how to improve their chance of reaching their targets are still understudied. Although a few *in vitro* studies provide valuable insights on how T cells migrate in 3D matrix structure, it is very difficult to track T cells within human tumor tissues(*15, 16*). Compared to normal tissues or any 3D matrix structures created in an in vitro experiment, ECM throughout tumors is highly dysregulated with substantial chemical and structural differences. Hence, *in vitro* experiments on T cells migration may not accurately reflect this crucial part of antitumor immune response in human TME.

With the spatial transcriptomics data, we developed a novel method to identify migration trails of T cells *in vivo*, under the assumption that T cells in the TME are gradually increasing their exhaustion levels. ReMiTT could offer an unbiased approach to investigate T cell migration in the tumor. When applied to multiple human cancer samples, we made the following observations: 1) several chemokines known to direct T cell migration exhibited an increasing trend along the trails; 2) Lag3/Galection-3 axis is a potential regulator for T cell exhaustion during migration; 3) the phenotype of the migrating T cells resembles previously reported T cell clusters with mixture of proliferative markers and Treg signatures.

Tumor-associated macrophage may play a positive or a negative role in anti-tumor immunity (*47*). For example, M1 macrophages are pro-inflammatory and tumor-resistant, while M2 macrophages are anti-inflammatory and pro-tumor. While it is not feasible to distinguish the two cell types without single cell resolution, we observed several signature genes of activated anti-tumor macrophage, including FOLR2 (*48*), PIM1 and SCL11A1 (*49*), were upregulated on the migration trails. These observations might suggest a positive role of macrophage in T cell migration in the tumor microenvironment. However, it should be noted that further evidence will be needed to prove the physical interactions between TAMs and the migrating T cells.

Why is it possible to observe 3-D migration trails on a thin-layer 2-D slide? Most of the trails were found along the edges of the hemorrhagic necrosis regions, which are expected to be spherical, instead of flat. Hence, it is possible that the ‘trails’ are derived from the intersection between a 2-D manifold and the cutting surface of the slide. The 2-D manifold might be the thin gaps between blocks of tissues in the 3-D space. If this is true, the picture of real T cell migration in the tumor might be much more complicated than previously thought, where T cells can choose different directions to move through.

There are several limitations to this work. First, the Visium spatial RNA-seq data is not at the single-cell level. With each spot having a diameter of 55 microns, it can cover 0-10 cells, depending on the spot’s location. Therefore, our analysis is based on the assumption that if multiple T cells are present in one spot, their exhaustion levels are similar. It will be interesting to implement this approach to data produced using the next-generation spatial transcriptomic technologies, such as Visium HD. Second, the sensitivity of this method hinges on high-level of T cell infiltration and large-span of T cell exhaustion markers, which may not be observed for all the tumor spatial transcriptomics samples. Finally, the scope of this work remains exploratory, and most conclusions in this work remain descriptive. Future experiments with *in vivo* tracing of T cell trajectories will be needed to test the validity of migration trails identified by ReMiTT. Despite these limitations, findings in this study might provide novel insights into T cell migration and exhaustion in the tumor microenvironment and facilitate the improvement of cancer immunotherapies.

## Materials and Methods

### T cell infiltrated spots and exhaustion score

We only include spots with T cells for the analysis in this study. We identify T cell infiltrated spots using T cell surface markers CD8A, CD8B, CD4, CD3D, CD3E, and CD3G. We assigned a T cell exhaustion score for each of the T cell spots based on the expression of exhaustion markers in these spots. T cell exhaustion markers used in this analysis are PDCD1, LAG3, HAVCR2, TIGIT, CTLA4, ENTPD1 and TOX. These genes have been shown to have increased expression among exhausted T cells in numerous studies (*10, 50–57*) and we use their sum expression as the level of exhaustion of T cells in this study. This sum expression could be correlated with the number of T cells in each spot. Thus, we performed linear regression with the dependent variable being the sum of gene expression of all these exhaustion markers in a spot and the independent variable being the sum of gene expression of all the T cell markers in the same spot, based on the assumption that spots with more T cells would have higher expression of T cell surface markers. Then, the T cell exhaustion score of each T cell spot is defined as the *residuals* of this linear model, reflecting how exhausted the T cells are in this spot, after controlling for the number of T cells in it. For convenience, we shift all the residuals to the right until the smallest exhaustion score is 0.

### Potential start and end nodes of trails

Assuming that activated T cells experience gradual exhaustion as they navigate through TME, the starting point of a migration trail would be a T cell node with a low exhaustion score, while the endpoint would be a node with a high exhaustion score. We ranked all T cell nodes based on their exhaustion scores and selected the first 33% of nodes with the lowest exhaustion scores as the potential start points and the last 33% with the highest exhaustion scores as the potential endpoints. With these potential start and endpoints, we identified potential migration trails with two methods.

### Minimum-spanning path

The first method of trail identification attempts to connect start points to endpoints using Dijkstra’s algorithm(*58*). In this method, we defined a directed, weighted graph G(V,E) with all the T cell nodes and their spatial locations. In this graph, V stands for vertices, in our case, all the T cell nodes. E stands for edges. The 10x Visium spatial gene expression data are organized in a hexagonal grid. Each spot (except those at the boundary of the slide) has 6 neighboring spots, with the distance between adjacent spots being approximately 110 microns. On graph G, edges only exist between neighboring nodes, with 2D distances smaller than 130 microns. In other words, we do not allow for spatial gaps along consecutive pairs along the migration trail. The direction of edges is set to point from the node with the smaller exhaustion score to the node with the larger exhaustion score. Then, for a pair of potential start and end node, if they can be connected on graph G, by a few intermediate spots, then we have a potential migration trail with T cells gradually getting more and more exhausted. It is possible that two nodes can be connected by multiple routes. Among these routes, we chose the one with gradual changes in T cell exhaustion stage rather than having big jumps in exhaustion stages between consecutive pairs. Thus, we assign the weight of edges to be the difference in exhaustion score between the two spots and use Dijkstra’s algorithm to find the shortest spanning path based on the weights of edges(*59, 60*). For each of the pairs of potential start and end nodes, we apply this method and got potential T cell migration trails. Then, we exclude trails featuring short distances between start and end nodes (< 500 microns) and trails comprising fewer than 5 nodes.

### Line in 3-D space

The minimum spanning path method, while helping to identify trails on which T cells have gradually changing exhaustion stages, requires exhaustion score to be strictly increasing along the trail. The exhaustion score in this study is a rough measure on T cell exhaustion stage. To require it to be strictly increasing along the way might exclude some of the real trails. Thus, we developed the 3-D line method to release this limitation and allow small fluctuations on exhaustion scores along the route while making sure the trend is still an increasing one.

For each T cell node, we use X and Y to represent its 2-dimensional coordinates on the tissue slide, and Z to represent its exhaustion score. Consequently, the 3-dimensional coordinates formed by X, Y, and Z of the T cell nodes along a migration trail would approximate a segment of a straight line in this 3-D space for those T cells migrating roughly in a linear pattern. To get this line segment, we first rescaled the XY coordinates of T cell nodes based on the minimum and maximum value of the exhaustion score (Z coordinate) in this data sample. After rescaling, the range of both X and Y coordinates becomes the same as the range of Z coordinates. Then, for each pair of potential start and end node, we construct a straight line in the 3D space connecting these two nodes. We then calculate the 3-D distance between all other T cell nodes and this 3-D line and keep the ones that have a very small distance to this line as a potential migration trail. The cutoff point is set arbitrarily but we advise users to use a small value. We then determine the indexing of nodes along the trail based on projections of each node to this straight line, beginning from the end with the smaller exhaustion scores.

Similar to the minimum spanning path method, we only include trails with at least 5 nodes as potential migration trails and exclude trails with start and end node too close to each other. Furthermore, we only kept trails with all the T cell nodes adjacent to each other (<=130 microns in 2D (XY) distance). Finally, while a rough line segment ensures that the trend of exhaustion score is increasing, small fluctuations is permissible. However, significant drawbacks between consecutive pairs of nodes in exhaustion score along the trail should be avoided so we removed such trails.

### Trail selection

With potential migration trails identified using these two strategies, we select the final set of trails based on the assumption that as a stream of T cells become more and more exhausted while migrating along some collagen fiber loosen area, the correlation of expression of T-cell stage markers of consecutive pairs of T cells on a migration trail would be higher compared to the correlation of the expression of these genes of T cells in neighboring areas. To make this comparison, we first obtained 5-15 alternative routes between the start and end point for each potential migration trail (Figure 1). Specifically, for a migration trail with n nodes, we first get n circular areas with a radius being 1000 microns, and the center being each node. Then, within the oval region surrounding the trail, formed by these overlapped circular areas, we obtained alternative routes of T cell nodes connecting the same pair of start and end nodes. For the alternative trails, there is no requirement for an increasing exhaustion score along the way, but all the other criteria for a potential migration trail apply, such as consecutive pairs of nodes along the trail must be neighbors and a trail must have >=6 spots and each spot should have T cell surface marker expression.

Among markers indicating T cell stages (*39*), we selected the ones that significantly positively correlated with the level of expression of T cell markers for this filtering. For each migration trail, we calculated the mean pairwise correlation of the normalized expression level of the selected T-cell stage-related genes(*61*). We then calculated the mean pairwise correlation for each of its alternative routes. Final set of migration trails all met two criteria: 1) mean pairwise correlation larger than all of its own alternative routes; 2) mean pairwise correlation larger than 85^th^ percentile of all alternative routes.

### Systematic analysis of algorithm-identified migration trails

#### 1) Test the trend of cytokine and chemokine receptors along the algorithm-identified migration trails

We assigned an index to each spot along the migration trails identified by our algorithm. Specifically, the starting spot was assigned an index of 1, with subsequent indices assigned sequentially along the trail until the endpoint. For CXCL9/10/11, we calculated the total expression of these ligands and performed a mixed linear regression analysis. In this model, the total expression of the ligands (normalized for UMI of each spot) served as the dependent variable, while the indices of spots along the migration trails were the independent variable. Migration trails were treated as random effects. A positive beta coefficient for the spot indices, accompanied by a p-value smaller than 0.05, indicates an increasing trend in the expression of these chemokines along the migration trail. Additionally, we visualized the gene expression levels of these chemokines at each index along the trail using boxplots. The same analysis was performed for CCL4/5 and CXCL16. These analyses were performed with the lung cancer sample, ovarian cancer sample, and melanoma sample.

Similarly, we tested the trend of markers of macrophages (total expression of CD68, CD163, CD80, CD14) (*62, 63*) with this method.

#### 2) Assign matched control trails for each algorithm-identified migration trails

First, for each of the algorithm-identified migration trails, we generated one matched control trail. To do this, we first performed clustering analysis with variable genes in the spatial data and divided the spots into six clusters based on dimensionality reduction methods(*64*). For a migration trail, one matched control trail was then generated with the *same length* as the migration trail and within the *same cluster* where the migration trail belongs to. Additional criteria for migration trail identification were also applied when generating the control trail, including requiring consecutive pairs along the trail to be adjacent to each other, all spots on the trail need to be T cell spots, and ensuring the spatial distance between the start and end spots of a trail to be ≥ 0.5 mm. For each of the *n* migration trails defined by the algorithm, we get one matched control trail and we call these *n* control trails one control set. We in total generated 5,000 matched control sets for each tumor sample.

#### 3) Shared TCR clones along trails compared to control trails

The test of whether T cells on migration trails are more likely to descend from the same clone compared to T cells not on these trails can only be performed for the first ovarian tumor sample, as the other samples do not have TCR variable genes. For each of the 19 migration trails identified in this sample, we calculated the mean number of shared TCR variable genes between consecutive pairs on the trails.

Specifically, for each trail, we examined the number of shared TCR variable genes between neighboring pairs of spots along the trail. The mean number of shared TCR variable genes for each trail is then calculated by averaging these values. The median number of shared TCR variable genes of the 19 migration trails are then compared to the empirical distribution calculated by the 5,000 control sets, and we reported the two-sided empirical p values. We performed the same analysis for BCR variable genes as B cells might as well migrate along these ‘physical highways’ and if this is the case, they can be sharing more BCR variables compared to those on control trails.

#### 4) Gene set enrichment analysis

We used transcripts per million (TPM) for gene expression in each spot to systematically analyze observed trails. To identify upregulated genes and gene pathways on these identified migration trails, we calculated the mean expression for each of the 3000 variable genes among migration trails and control sets. We then calculated empirical p values for each gene using the empirical distribution generated by the 5,000 control sets. Benjamini-Hochberg adjustment was performed on these p-values to account for multiple comparisons. With genes significantly overexpressed on migration trails compared to the control sets, we performed gene set enrichment analysis (GESA)(*38*). We used C5 collection GO term of Human Molecular Signature Dataset (MSigDB) for this analysis (*65*). Go term contains 10402 gene pathway sets that are related to oncology.

#### 5) Phenotype of migrating T cells

We then characterized the phenotype of the moving T cells on our migration trails using deep single-cell RNA-seq data from 12,346 T cells from 14 non-small cell lung cancer patients(*39*). Initially, we performed clustering analysis and divided these T cells into 13 clusters(*64*). Next, we calculated the mean expression for each of the significantly upregulated genes on our migration trails within each cluster. For each gene, we ranked the 13 means from smallest to largest, with rank 13 corresponding to the largest mean among the clusters. The T cell clusters with more top ranks indicate a higher enrichment of markers from our migration trails, suggesting that these clusters represent the phenotype of the migrating T cells on the identified trails.

## Supplementary Figures

**Fig S1.**
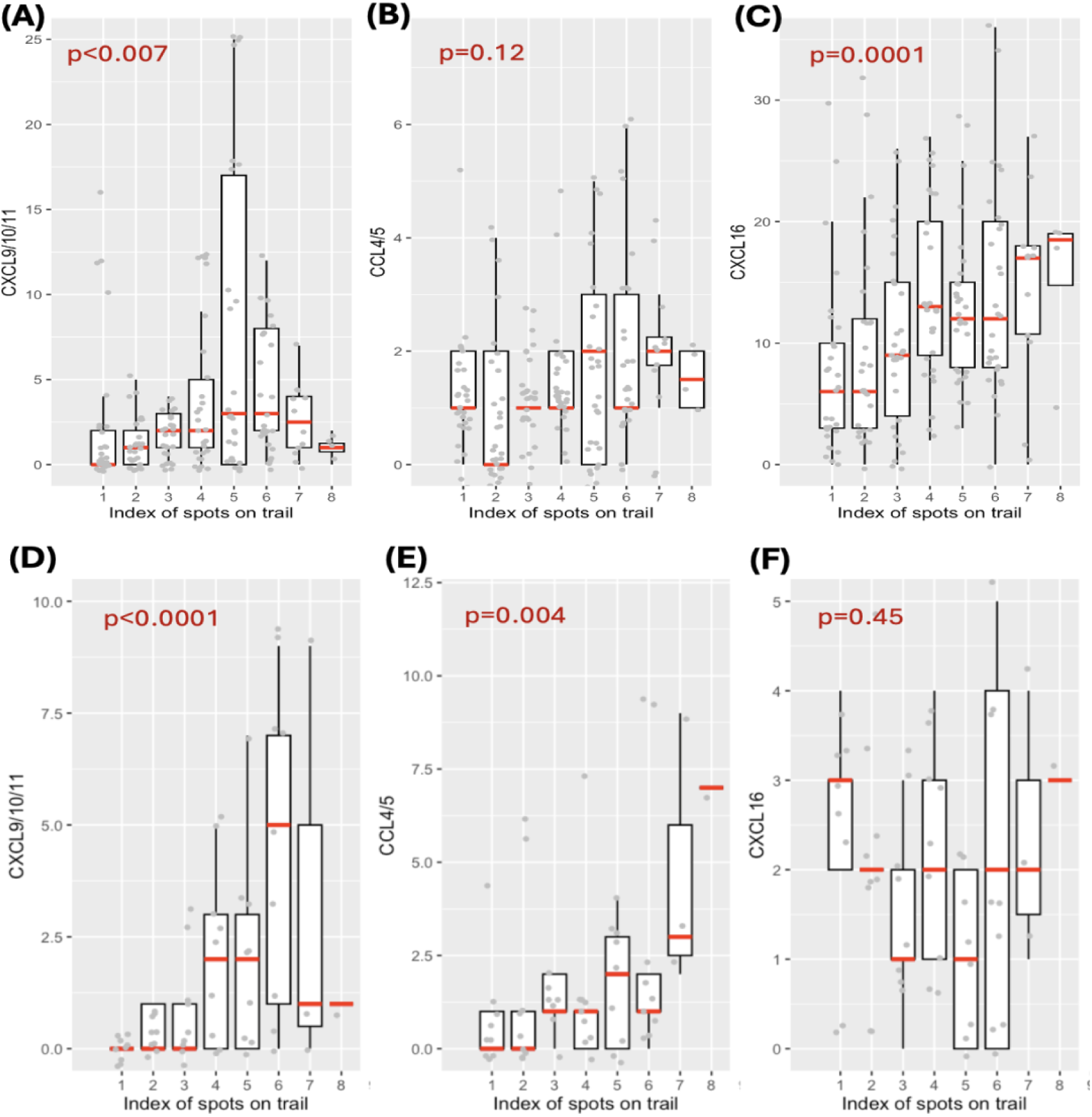
Trend of chemokines that promote T cell migration along the algorithm-identified migration trails, and phenotype of these trails identified in the melanoma sample. **(A)-(C)**Total expression of CXCL9, CXCL10, and CXCL11, Total expression and CCL4 and CCL5 and expression of CXCL16 along the algorithm identified migration trails in the **ovarian cancer sample** with the median of each index labeled with a red line segment. Mixed-effect linear regression adjusting for UMI of each spot is performed to test the increasing trend of ligand expression along the trails, and 2-sided p value is reported in the panel. **(D)-(F)** Total expression of CXCL9, CXCL10, and CXCL11, Total expression and CCL4 and CCL5 and expression of CXCL16 along the algorithm identified migration trails in the **melanoma sample** with the median of each index labeled with a red line segment. Mixed-effect linear regression adjusting for UMI of each spot is performed to test the increasing trend of ligand expression along the trails, and 2-sided p value is reported in the panel.

**Fig. S2.**
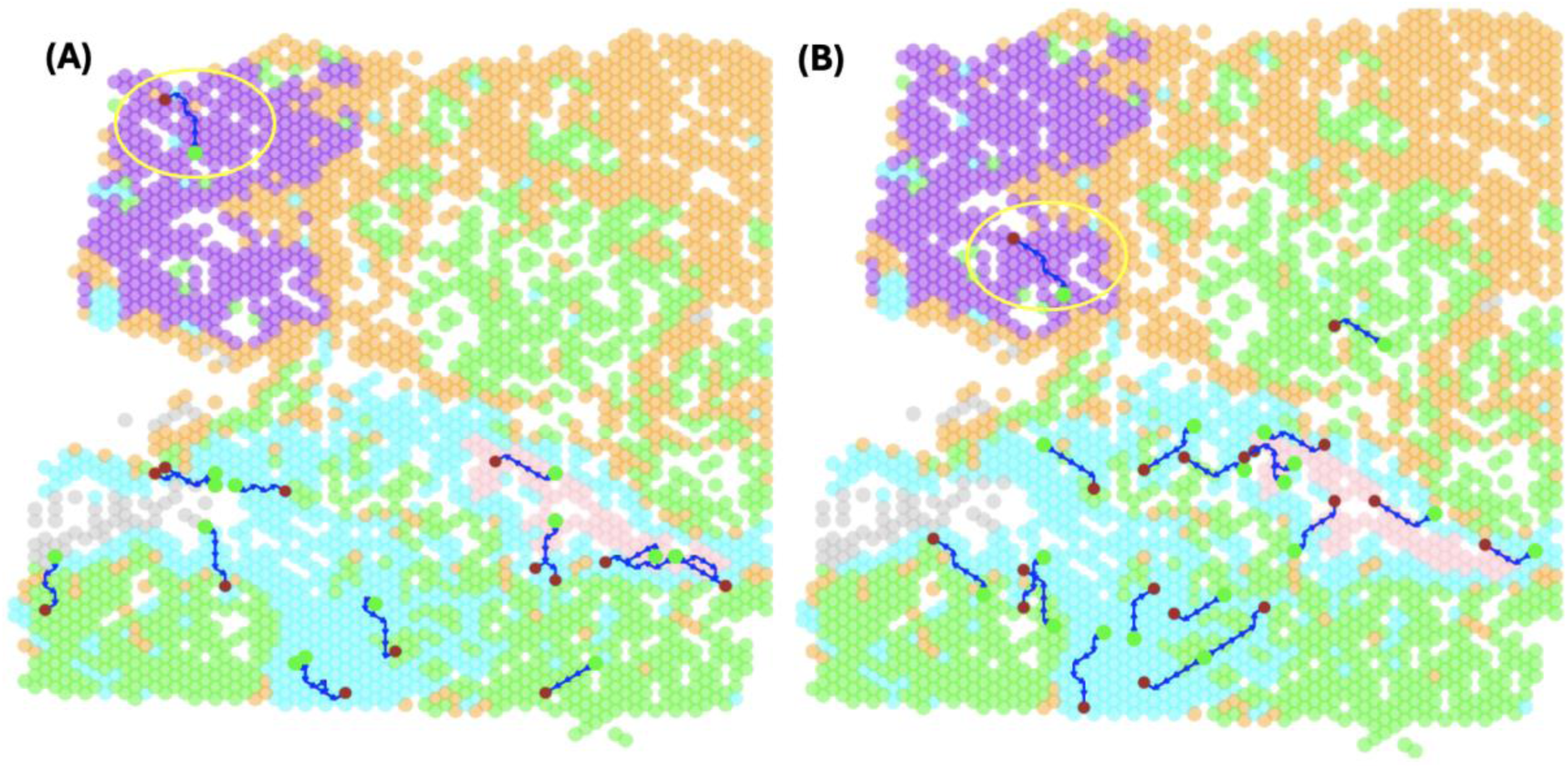
An example of a matched control set paired with algorithm-identified migration trails. **(A)**The 19 algorithm-identified migration trails. **(B)** One matched control set-matched trails corresponding to each of the migration trails. Colored spots represent different gene-expression clusters. On each trail, the green spot marks the start point, and the brown spot indicates the end point. Each algorithm-identified trail is paired with a control trail of the same gene-expression cluster and length. All control trails are generated with T cell spots only, same as the migration trails. The circled pair of trails shows an example of an algorithm-identified trail and its corresponding control trail. While (B) shows one control trail set, a total of 5,000 control sets were generated for systematic analysis.

**Fig. S3.**
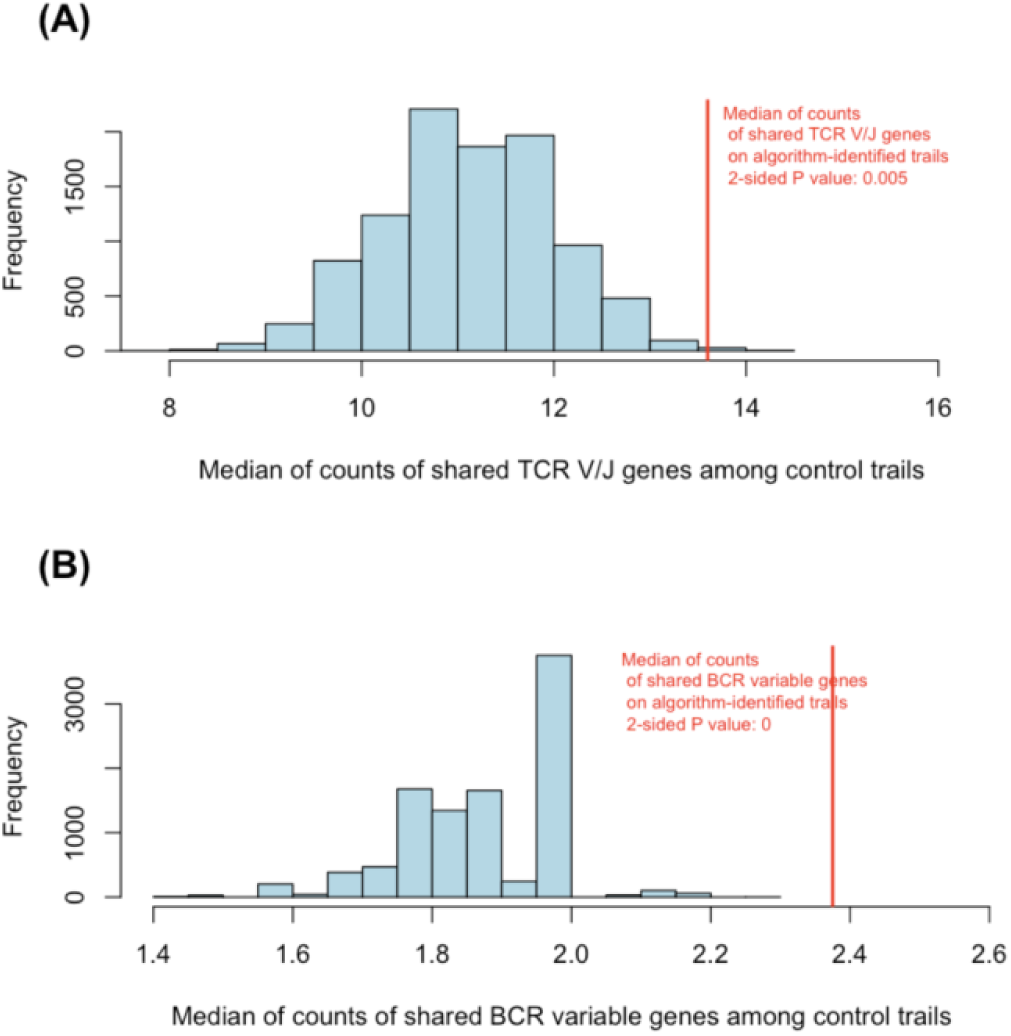
Comparison of shared TCR and BCR variable genes between algorithm-identified trails and control trails in an ovarian tumor sample with TCR and BCR variable genes. **(A)**The histogram shows the empirical distribution of the median of mean counts of shared TCR V/J genes between consecutive pairs within each control trail in a 25-trail set for 5,000 such control trail sets. The red line indicates the median of mean counts of shared TCR V/J genes for the 25 algorithm-defined migration trails (median = 13.6). The two-sided p-value for this comparison is 0.005. **(B)** The histogram shows the empirical distribution of the median of mean counts of shared BCR variable genes between consecutive pairs within each control trail in a 25-trail set for 5,000 such control trail sets. The red line marks the median count of shared BCR variable genes for the algorithm-defined migration trails (median = 2.375). The two-sided p-value for this comparison is <0.0001

**Fig S4.**
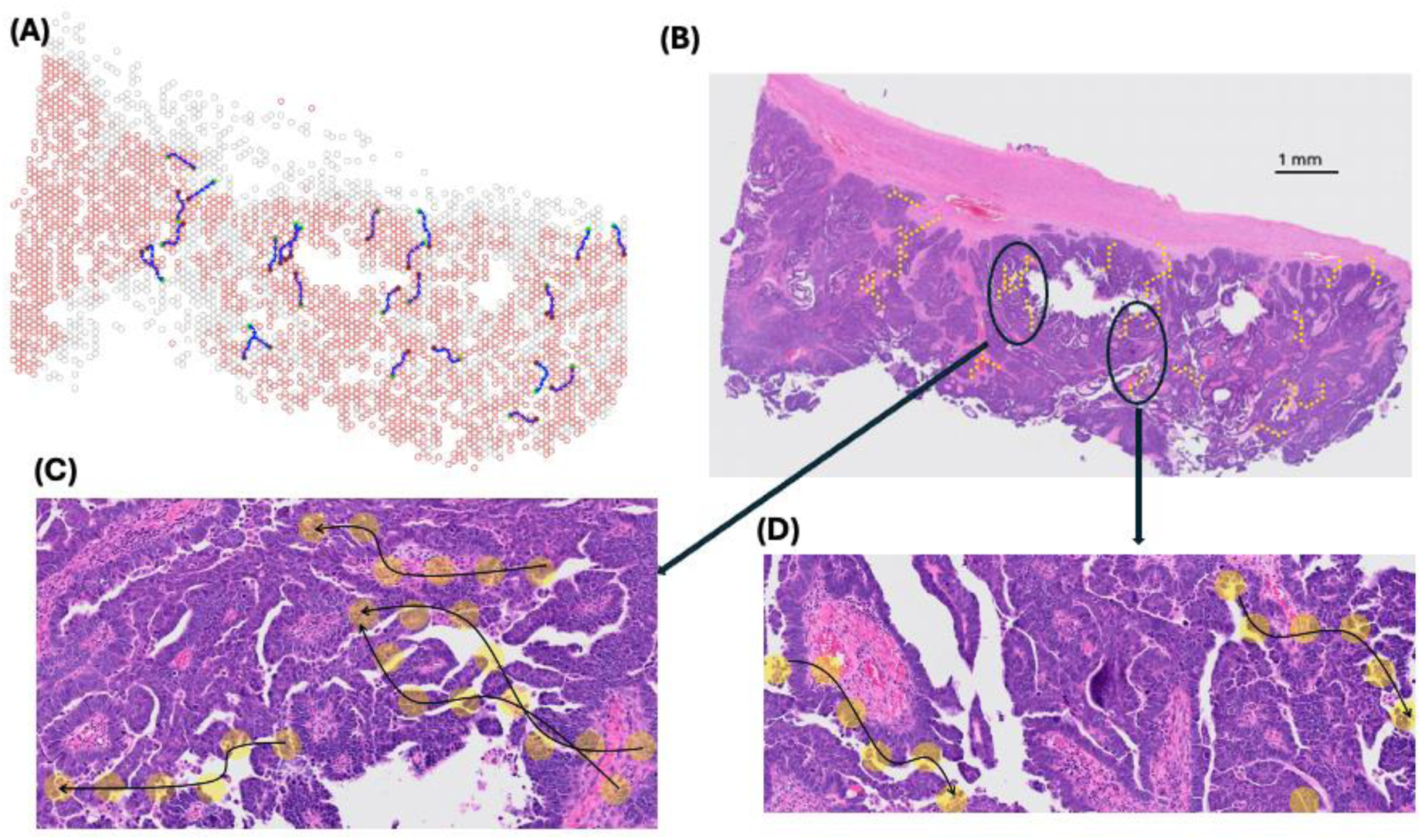
Migration trails identified in a human ovarian cancer sample and their projections on the pathology slide. **(A)**Algorithm-identified migration trails labeled as splines on the spatial coordinates, with the starting location marked as green. Grey spots are spots with T cell infiltration and red circles indicating spots with high levels of expression of ovarian cancer markers. **(B)** Same trails overlaid on the pathology (H&E) slide of the tumor. **(C)** Examples of trails travel in less dense TME. **(D)** Examples of trails whose origin is close to vessels.

**Fig. S5.**
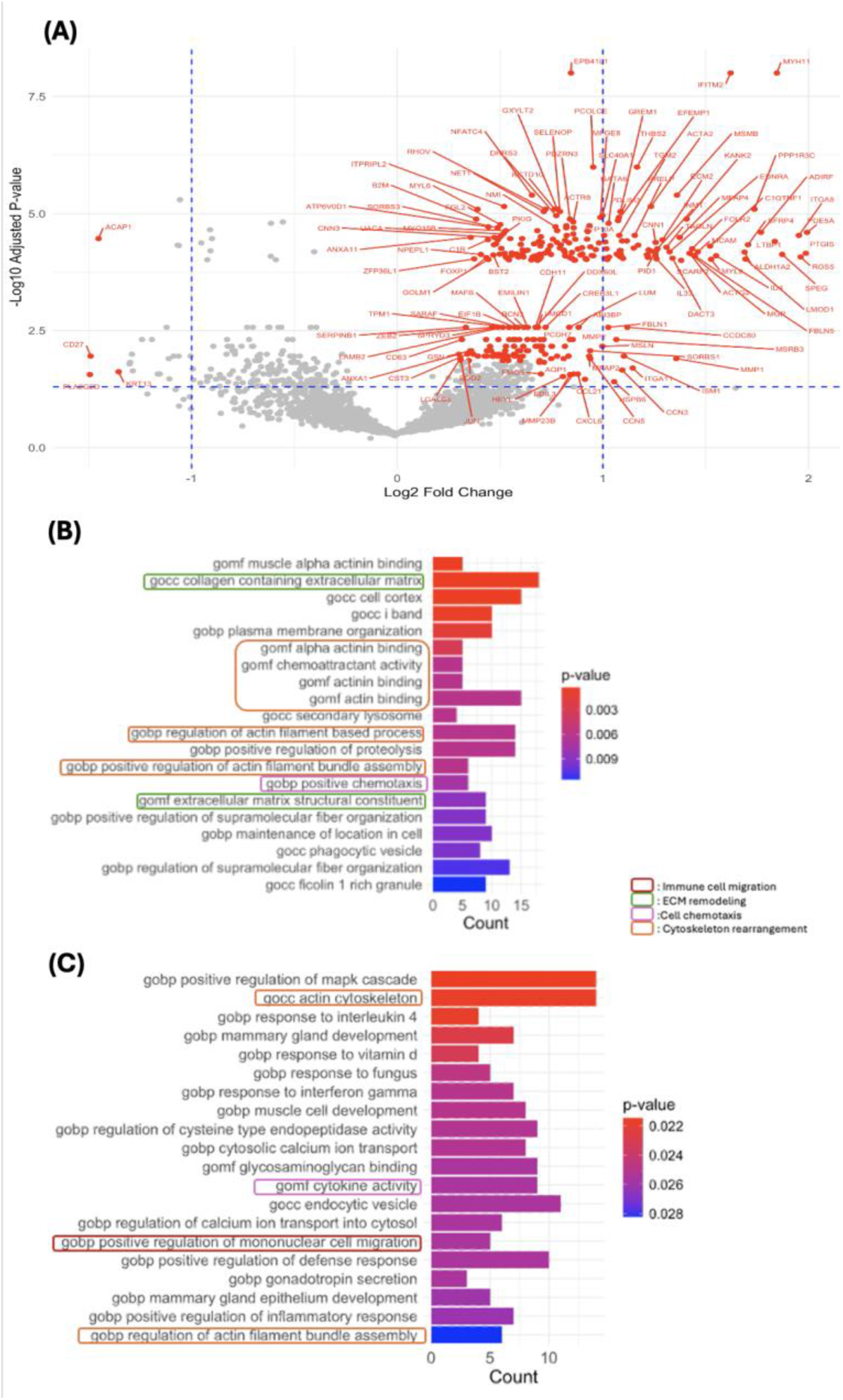
Differential gene expression analysis for T cell migration trails in the ovarian cancer sample. **(A)** Volcano plot of gene (3000 variable genes) fold change (FC) and adjusted empirical p values between those on migration trails vs. 5000 control sets. Genes with FDR<0.05 and the absolute values of FC ≥1.2 were labeled on the plot. **(B)** Barplot showing the top 20 enriched Gene Ontology biological process (GOBP) gene pathways computed from the 191 significantly upregulated genes on migration trails by GSEA. Colors of the textbox borders indicated the type of biological processes that may involved in T cell migration. Colors of the bars indicate the adjusted p values. Statistical significance was evaluated using Fisher’s exact test, with FDR corrected by the Benjamini-Hochberg approach. **(C)** Barplot showing the top 40-60 enriched Gene Ontology biological process (GOBP) gene pathways computed from the 191 significantly upregulated genes on migration trails by GSEA.

**Fig S6.**
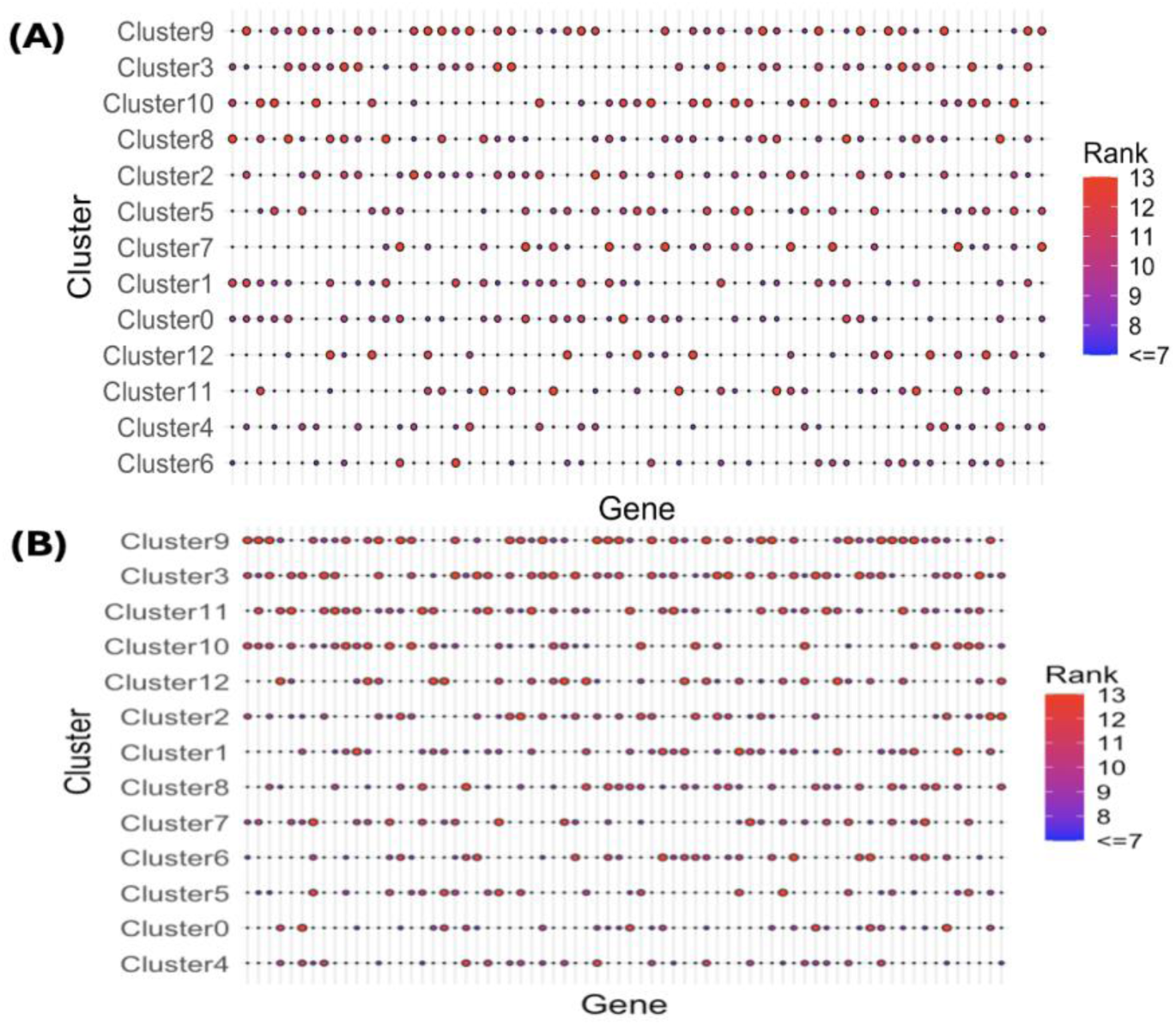
Enrichment of overexpressed genes on T cell migration trails in each of the T cell clusters in the Lung single-cell cohort for ovarian cancer sample and melanoma sample. **(A)**Bubble plot showing the enrichment of overexpressed genes on T cell migration trails in each of the T cell clusters in Figure 5(A) in the ovarian cancer sample. The large bubble sizes and red colors indicate higher ranks of mean expression of each of the upregulated genes on migration trails among all the 13 clusters of T cells. Clusters (y-axis) were ordered by their median ranks of the genes. **(B)** Bubble plot showing the enrichment of overexpressed genes on T cell migration trails in each of the T cell clusters in Figure 5(A) for the melanoma sample.

**Fig. S7.**
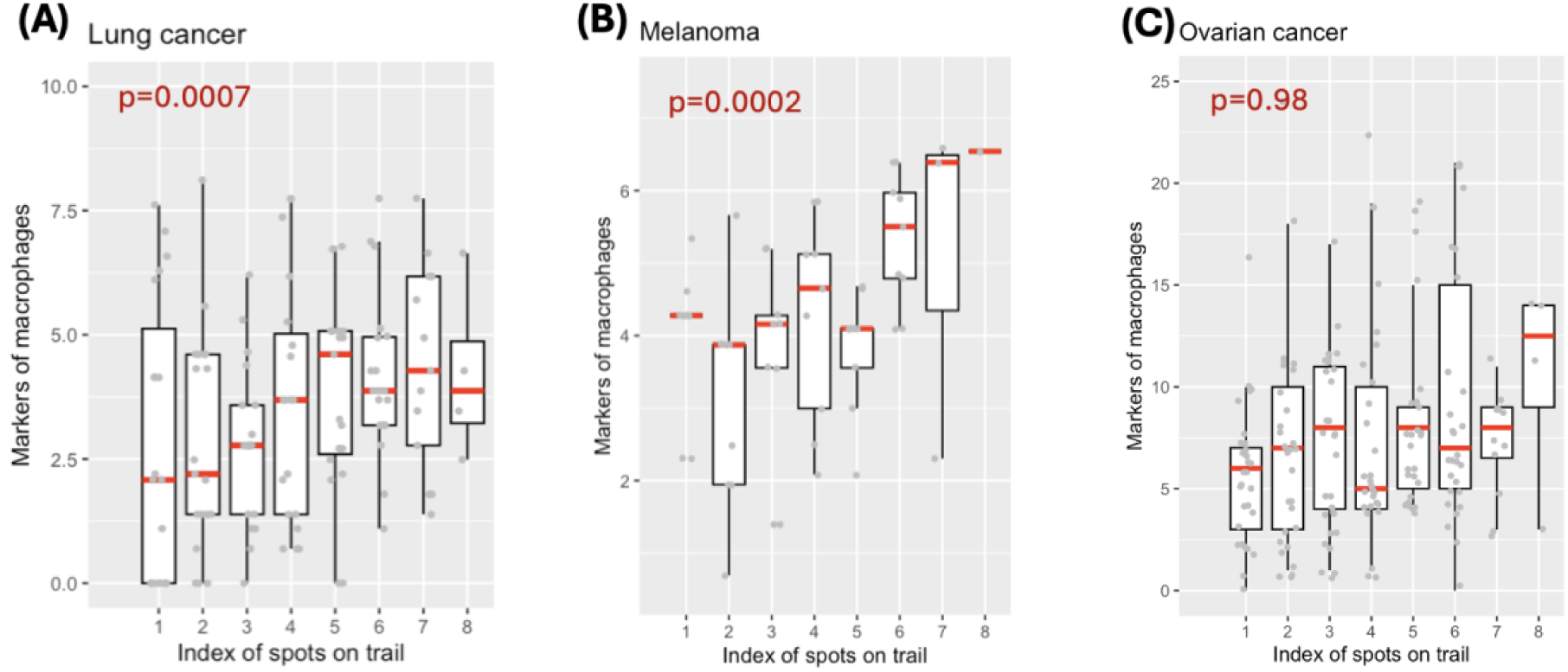
Trend of the expression of macrophage markers along the algorithm-identified migration trails. Total expression of macrophage markers (CD68, CD163, CD80, CD14) along the algorithm identified migration trails for the sample of lung cancer **(A),** melanoma **(B),** and ovarian cancer **(C),** with the median of each index labeled with a red line segment. Mixed-effect linear regression adjusting for UMI of each spot is performed to test the increasing trend of the expression along the trails, and 2-sided p values are reported in the panel.

## Author contributions

Conceptualization: LZ, BL, QL, GX Methodology: LZ, QL, GX Investigation: LZ, BL, QL, GX,SZ Visualization: LZ, QL, GX Supervision: QL, GX Writing—original draft: LZ, BL,QL, GX, SZ Writing—review & editing: LZ, BL,QL, GX, SZ

## Data and materials availability

The authors analyzed two publicly available SRT data. Raw count matrices, images, and spatial information for two SRT data from 10x Visium are accessible on the 10x Genomics website at https://support.10xgenomics.com/spatial-gene-expression/datasets. The first sample can be downloaded at https://www.10xgenomics.com/datasets/human-ovarian-cancer-1-standard and the second sample can be downloaded at https://www.10xgenomics.com/datasets/human-ovarian-cancer-11-mm-capture-area-ffpe-2-standard. The deep sequencing single cell data for T cells can be downloaded from: https://www.ncbi.nlm.nih.gov/geo/query/acc.cgi?acc=GSE99254

An open-source implementation of the algorithm in R is available at: https://github.com/zhonglin1234/ReMiTT

