## Supplementary tables for "Computational Identification of Migrating T cells in Spatial Transcriptomics Data"

**Table S1. Upregulated genes on the migration trails in the lung sample**

|  | **Fold change** | **Adjusted p values** |
| --- | --- | --- |
| PLN | 9.08 | 0.000 |
| SNCG | 7.01 | 0.000 |
| CCDC3 | 6.50 | 0.000 |
| OGN | 5.52 | 0.000 |
| COL14A1 | 4.00 | 0.000 |
| PDE5A | 3.98 | 0.000 |
| PTGIS | 3.96 | 0.000 |
| RGS5 | 3.89 | 0.000 |
| ITGA8 | 3.87 | 0.000 |
| SPEG | 3.66 | 0.000 |
| MYH11 | 3.60 | 0.000 |
| LTBP1 | 3.57 | 0.000 |
| ADIRF | 3.41 | 0.000 |
| PPP1R3C | 3.33 | 0.000 |
| SFRP4 | 3.26 | 0.000 |
| LMOD1 | 3.23 | 0.000 |
| ALDH1A2 | 3.22 | 0.000 |
| IFITM2 | 3.08 | 0.000 |
| ID4 | 2.93 | 0.000 |
| MYL9 | 2.88 | 0.000 |
| C1QTNF1 | 2.87 | 0.000 |
| MCAM | 2.75 | 0.000 |
| FBLN5 | 2.72 | 0.000 |
| FOLR2 | 2.70 | 0.000 |
| MGP | 2.68 | 0.000 |
| ACTG2 | 2.68 | 0.000 |
| KANK2 | 2.65 | 0.000 |
| SCARF2 | 2.60 | 0.000 |
| MFAP4 | 2.59 | 0.000 |
| MSMB | 2.57 | 0.000 |
| MMP1 | 2.56 | 0.013 |
| DACT3 | 2.53 | 0.000 |
| EDNRA | 2.50 | 0.000 |
| TAGLN | 2.48 | 0.000 |
| ECM2 | 2.45 | 0.000 |
| INMT | 2.40 | 0.000 |
| IL33 | 2.40 | 0.000 |
| PID1 | 2.39 | 0.000 |
| CNN1 | 2.39 | 0.000 |
| CXCL12 | 2.39 | 0.000 |
| IGFBP7 | 2.35 | 0.000 |
| ACTA2 | 2.35 | 0.000 |
| IER5L | 2.33 | 0.000 |
| C11orf96 | 2.33 | 0.000 |
| PDGFRA | 2.33 | 0.000 |
| LHFPL6 | 2.32 | 0.000 |
| MRVI1 | 2.25 | 0.000 |
| THBS2 | 2.24 | 0.000 |
| PRELP | 2.23 | 0.000 |
| PPP1R14A | 2.22 | 0.000 |
| ITGA11 | 2.21 | 0.020 |
| CCDC80 | 2.17 | 0.003 |
| FOXC1 | 2.17 | 0.000 |
| ISM1 | 2.15 | 0.011 |
| CPED1 | 2.14 | 0.000 |
| CCN3 | 2.13 | 0.022 |
| RASL12 | 2.13 | 0.000 |
| FCGBP | 2.13 | 0.000 |
| ADAMTS2 | 2.12 | 0.000 |
| CD28 | 2.12 | 0.000 |
| GREM1 | 2.12 | 0.000 |
| BGN | 2.12 | 0.000 |
| TGM2 | 2.11 | 0.000 |
| PDLIM3 | 2.11 | 0.000 |
| CD34 | 2.11 | 0.000 |
| EFEMP1 | 2.10 | 0.000 |
| RGL3 | 2.10 | 0.000 |
| MSRB3 | 2.09 | 0.005 |
| HSPB6 | 2.08 | 0.039 |
| FBLIM1 | 2.04 | 0.000 |
| GATA6 | 2.04 | 0.000 |
| FBLN1 | 2.04 | 0.003 |
| DEPP1 | 2.03 | 0.000 |
| TMEM204 | 2.02 | 0.000 |
| EVA1B | 2.02 | 0.000 |
| CAMK1 | 2.01 | 0.000 |
| THY1 | 2.01 | 0.000 |
| SLC40A1 | 2.00 | 0.000 |
| MSLN | 2.00 | 0.007 |
| MFGE8 | 1.98 | 0.000 |
| SORBS2 | 1.97 | 0.000 |
| MUC5B | 1.95 | 0.000 |
| AEBP1 | 1.94 | 0.000 |
| PCOLCE | 1.94 | 0.000 |
| SFRP2 | 1.93 | 0.000 |
| CCN2 | 1.92 | 0.000 |
| CCN5 | 1.92 | 0.013 |
| SORBS1 | 1.92 | 0.008 |
| ATP10A | 1.91 | 0.000 |
| MFAP2 | 1.91 | 0.011 |
| GUCY1A1 | 1.90 | 0.000 |
| SH3RF1 | 1.90 | 0.000 |
| A2M | 1.89 | 0.000 |
| CEACAM6 | 1.89 | 0.000 |
| FMO2 | 1.89 | 0.000 |
| CCL21 | 1.88 | 0.035 |
| OLFML2B | 1.87 | 0.000 |
| ATP6V1H | 1.86 | 0.000 |
| GFPT2 | 1.85 | 0.000 |
| ABI3BP | 1.84 | 0.003 |
| CXCL6 | 1.84 | 0.027 |
| IFITM3 | 1.84 | 0.000 |
| KIF1C | 1.82 | 0.000 |
| PRRX1 | 1.82 | 0.000 |
| METRN | 1.82 | 0.000 |
| MMP23B | 1.81 | 0.027 |
| FAM174B | 1.81 | 0.000 |
| PRUNE2 | 1.81 | 0.000 |
| ACTR8 | 1.81 | 0.000 |
| MMP2 | 1.81 | 0.005 |
| LDOC1 | 1.80 | 0.000 |
| JCAD | 1.80 | 0.000 |
| EPB41L1 | 1.80 | 0.000 |
| ANTXR1 | 1.79 | 0.000 |
| EDIL3 | 1.79 | 0.027 |
| PCDH7 | 1.79 | 0.005 |
| PDZRN3 | 1.79 | 0.000 |
| LUM | 1.79 | 0.003 |
| ARHGEF17 | 1.78 | 0.000 |
| COL6A2 | 1.78 | 0.000 |
| CLEC14A | 1.78 | 0.011 |
| PTGS2 | 1.78 | 0.011 |
| KALRN | 1.77 | 0.000 |
| CMPK2 | 1.77 | 0.000 |
| MYLK | 1.77 | 0.000 |
| SCGB1A1 | 1.76 | 0.000 |
| RHOB | 1.76 | 0.000 |
| CMKLR1 | 1.76 | 0.007 |
| CALD1 | 1.76 | 0.000 |
| CAVIN1 | 1.75 | 0.000 |
| AQP1 | 1.75 | 0.013 |
| HEYL | 1.75 | 0.030 |
| FSTL1 | 1.73 | 0.000 |
| SELENOP | 1.73 | 0.000 |
| TIMP2 | 1.73 | 0.000 |
| GXYLT2 | 1.71 | 0.000 |
| FLNA | 1.71 | 0.000 |
| KLHDC8B | 1.71 | 0.011 |
| COL1A2 | 1.71 | 0.000 |
| DHRS3 | 1.71 | 0.000 |
| NMI | 1.70 | 0.000 |
| KCTD10 | 1.70 | 0.000 |
| SPARCL1 | 1.69 | 0.000 |
| MGMT | 1.68 | 0.007 |
| EFEMP2 | 1.68 | 0.007 |
| HCFC2 | 1.68 | 0.013 |
| GLI3 | 1.68 | 0.000 |
| COL6A1 | 1.68 | 0.000 |
| NR1D1 | 1.66 | 0.000 |
| CREB3L1 | 1.65 | 0.003 |
| RHOV | 1.65 | 0.000 |
| NRP1 | 1.65 | 0.000 |
| CD248 | 1.65 | 0.010 |
| PRDM6 | 1.64 | 0.015 |
| NET1 | 1.64 | 0.000 |
| CPE | 1.64 | 0.005 |
| SPARC | 1.63 | 0.007 |
| SERPING1 | 1.63 | 0.000 |
| TJP3 | 1.63 | 0.008 |
| SLC4A4 | 1.62 | 0.000 |
| FMO5 | 1.62 | 0.027 |
| C1S | 1.62 | 0.000 |
| NNMT | 1.62 | 0.000 |
| LRP1 | 1.62 | 0.000 |
| COL1A1 | 1.62 | 0.005 |
| MXRA8 | 1.62 | 0.000 |
| SH3BGRL | 1.62 | 0.007 |
| SOD3 | 1.62 | 0.014 |
| ETS2 | 1.61 | 0.000 |
| RARRES2 | 1.61 | 0.015 |
| TIMP1 | 1.61 | 0.000 |
| CDS1 | 1.61 | 0.000 |
| LMCD1 | 1.61 | 0.003 |
| PPIA | 1.60 | 0.000 |
| LIX1L | 1.60 | 0.000 |
| CD151 | 1.60 | 0.000 |
| LIFR | 1.60 | 0.007 |
| ARRDC2 | 1.60 | 0.007 |
| DDX60L | 1.60 | 0.003 |
| CLU | 1.60 | 0.000 |
| MAP3K8 | 1.59 | 0.007 |
| CDH11 | 1.59 | 0.003 |
| VCAN | 1.59 | 0.000 |
| SYTL3 | 1.59 | 0.005 |
| NBL1 | 1.59 | 0.000 |
| FCER1G | 1.58 | 0.005 |
| ETS1 | 1.58 | 0.000 |
| GNB4 | 1.57 | 0.000 |
| TGFB1I1 | 1.57 | 0.035 |
| NFATC4 | 1.57 | 0.000 |
| CBX7 | 1.57 | 0.000 |
| YPEL2 | 1.56 | 0.018 |
| AKAP12 | 1.56 | 0.020 |
| ELN | 1.56 | 0.021 |
| COL16A1 | 1.55 | 0.008 |
| EMILIN1 | 1.55 | 0.003 |
| TNS2 | 1.55 | 0.026 |
| PPP1R12B | 1.55 | 0.000 |
| GLT8D1 | 1.55 | 0.005 |
| RRAS | 1.55 | 0.008 |
| JDP2 | 1.55 | 0.008 |
| PLEKHH2 | 1.54 | 0.033 |
| FILIP1L | 1.54 | 0.015 |
| SULF1 | 1.54 | 0.026 |
| ARMCX3 | 1.54 | 0.035 |
| RCN3 | 1.53 | 0.003 |
| CD81 | 1.53 | 0.000 |
| DCLK2 | 1.53 | 0.035 |
| DSTN | 1.53 | 0.000 |
| VAMP5 | 1.53 | 0.007 |
| SYNPO | 1.53 | 0.011 |
| LAMB1 | 1.53 | 0.000 |
| HOXB4 | 1.52 | 0.022 |
| VIM | 1.52 | 0.000 |
| DPYSL3 | 1.52 | 0.000 |
| HTRA1 | 1.52 | 0.000 |
| KCNMA1 | 1.52 | 0.018 |
| FBN1 | 1.51 | 0.014 |
| BPIFB1 | 1.51 | 0.014 |
| PPP2CB | 1.51 | 0.008 |
| ABCA6 | 1.51 | 0.033 |
| SNX21 | 1.51 | 0.005 |
| MTUS1 | 1.51 | 0.008 |
| SDCBP | 1.51 | 0.016 |
| NFIA | 1.51 | 0.008 |
| PRKG1 | 1.50 | 0.008 |
| GNAI2 | 1.50 | 0.000 |
| MAFB | 1.50 | 0.003 |
| AKAP6 | 1.50 | 0.016 |
| DNAH5 | 1.50 | 0.037 |
| FAS | 1.50 | 0.028 |
| ANKDD1B | 1.50 | 0.011 |
| PDGFRB | 1.50 | 0.000 |
| COL3A1 | 1.50 | 0.025 |
| ZBTB47 | 1.49 | 0.041 |
| BIN3 | 1.49 | 0.005 |
| EIF1B | 1.49 | 0.003 |
| ADGRL2 | 1.48 | 0.011 |
| GLI2 | 1.48 | 0.010 |
| PDLIM7 | 1.48 | 0.007 |
| RHOU | 1.48 | 0.037 |
| DCN | 1.48 | 0.005 |
| C5AR1 | 1.47 | 0.019 |
| TPM2 | 1.47 | 0.000 |
| GPD1L | 1.47 | 0.045 |
| TNFSF12 | 1.47 | 0.027 |
| PER1 | 1.47 | 0.000 |
| SELENBP1 | 1.46 | 0.027 |
| WBP1L | 1.46 | 0.000 |
| ARHGEF25 | 1.46 | 0.041 |
| LYN | 1.46 | 0.005 |
| SARAF | 1.46 | 0.003 |
| LRTOMT | 1.46 | 0.023 |
| TNS1 | 1.46 | 0.000 |
| TPM1 | 1.46 | 0.003 |
| VCL | 1.45 | 0.000 |
| FMOD | 1.45 | 0.031 |
| ARHGEF10 | 1.45 | 0.014 |
| ACSF2 | 1.45 | 0.011 |
| FCGR2A | 1.45 | 0.013 |
| PARVA | 1.45 | 0.000 |
| CRISPLD2 | 1.45 | 0.041 |
| ISG15 | 1.44 | 0.013 |
| TAMM41 | 1.44 | 0.039 |
| TMEM43 | 1.44 | 0.007 |
| LTBP3 | 1.44 | 0.000 |
| MYO15B | 1.44 | 0.000 |
| HEMK1 | 1.44 | 0.033 |
| MYADM | 1.44 | 0.041 |
| FKBP10 | 1.44 | 0.018 |
| SMTN | 1.44 | 0.008 |
| NPDC1 | 1.44 | 0.023 |
| NME2 | 1.44 | 0.000 |
| APOL3 | 1.44 | 0.042 |
| C1QB | 1.43 | 0.010 |
| ITPRIPL2 | 1.43 | 0.000 |
| UACA | 1.43 | 0.000 |
| LAMB2 | 1.43 | 0.003 |
| IGFBP5 | 1.43 | 0.014 |
| AMDHD2 | 1.42 | 0.011 |
| FAM83E | 1.42 | 0.037 |
| PIGR | 1.42 | 0.021 |
| FLCN | 1.42 | 0.000 |
| ANGPTL2 | 1.42 | 0.026 |
| PKIG | 1.42 | 0.000 |
| FGL2 | 1.42 | 0.000 |
| ARHGEF40 | 1.41 | 0.047 |
| PMP22 | 1.41 | 0.039 |
| MYO1E | 1.41 | 0.000 |
| PLPP1 | 1.41 | 0.048 |
| CCND3 | 1.41 | 0.019 |
| CCDC102B | 1.41 | 0.028 |
| C1QC | 1.41 | 0.016 |
| DUSP1 | 1.41 | 0.010 |
| PROM1 | 1.41 | 0.011 |
| SPON2 | 1.41 | 0.028 |
| EGR1 | 1.41 | 0.005 |
| LXN | 1.40 | 0.019 |
| SPRYD3 | 1.40 | 0.003 |
| ACTN1 | 1.40 | 0.000 |
| FAM210B | 1.40 | 0.029 |
| STMN3 | 1.40 | 0.036 |
| SORBS3 | 1.40 | 0.000 |
| SDC3 | 1.40 | 0.030 |
| IL10RA | 1.40 | 0.011 |
| NUAK1 | 1.40 | 0.046 |
| ZNF703 | 1.40 | 0.000 |
| MAP2K3 | 1.39 | 0.010 |
| EPC1 | 1.39 | 0.040 |
| GEM | 1.39 | 0.016 |
| ZNF440 | 1.39 | 0.041 |
| CNN3 | 1.39 | 0.000 |
| IFI6 | 1.39 | 0.000 |
| IFT172 | 1.38 | 0.011 |
| HAGHL | 1.38 | 0.027 |
| BST2 | 1.38 | 0.000 |
| TSPAN4 | 1.38 | 0.039 |
| MYH10 | 1.38 | 0.011 |
| CELF2 | 1.38 | 0.011 |
| ZEB2 | 1.38 | 0.003 |
| C8orf34 | 1.37 | 0.042 |
| C3 | 1.37 | 0.031 |
| NFIX | 1.37 | 0.008 |
| ECM1 | 1.37 | 0.022 |
| ARVCF | 1.37 | 0.028 |
| IFI44L | 1.37 | 0.037 |
| GAS6 | 1.37 | 0.007 |
| SERPINH1 | 1.37 | 0.008 |
| IL18BP | 1.36 | 0.028 |
| GOLM1 | 1.36 | 0.000 |
| ATP6V0D1 | 1.36 | 0.000 |
| C1R | 1.36 | 0.000 |
| LRRC56 | 1.36 | 0.044 |
| DDAH2 | 1.35 | 0.008 |
| IFI35 | 1.35 | 0.008 |
| ETV3 | 1.35 | 0.021 |
| SLC9A3R2 | 1.35 | 0.011 |
| FOXP1 | 1.35 | 0.000 |
| TNKS1BP1 | 1.35 | 0.005 |
| LGALS1 | 1.34 | 0.035 |
| MT2A | 1.34 | 0.011 |
| PLBD2 | 1.34 | 0.039 |
| AXL | 1.34 | 0.036 |
| RBPMS | 1.34 | 0.027 |
| CSRP1 | 1.33 | 0.027 |
| PRKCE | 1.33 | 0.038 |
| IGFBP3 | 1.33 | 0.033 |
| NPEPL1 | 1.33 | 0.000 |
| ARHGAP1 | 1.32 | 0.032 |
| SULF2 | 1.32 | 0.030 |
| HSPG2 | 1.31 | 0.035 |
| UCP2 | 1.31 | 0.011 |
| DBP | 1.31 | 0.029 |
| PLAAT4 | 1.31 | 0.018 |
| MYL6 | 1.31 | 0.000 |
| TPP1 | 1.31 | 0.032 |
| TCP11L2 | 1.31 | 0.046 |
| STARD10 | 1.31 | 0.038 |
| B2M | 1.30 | 0.000 |
| METTL7A | 1.30 | 0.035 |
| NDUFAF3 | 1.30 | 0.018 |
| KLF6 | 1.30 | 0.034 |
| CFB | 1.30 | 0.039 |
| ZFP36L1 | 1.30 | 0.000 |
| HMGN3 | 1.29 | 0.010 |
| ANKRD65 | 1.29 | 0.028 |
| PNPLA2 | 1.29 | 0.037 |
| S100A11 | 1.28 | 0.008 |
| CCDC85B | 1.28 | 0.035 |
| ANXA11 | 1.28 | 0.000 |
| CD74 | 1.28 | 0.024 |
| GSN | 1.28 | 0.008 |
| EPS15 | 1.27 | 0.020 |
| COL4A1 | 1.27 | 0.025 |
| SOD2 | 1.27 | 0.014 |
| UNC93B1 | 1.27 | 0.041 |
| GAS7 | 1.27 | 0.037 |
| ZFP36L2 | 1.27 | 0.021 |
| CST3 | 1.27 | 0.010 |
| CHKA | 1.27 | 0.049 |
| SERPINB1 | 1.26 | 0.003 |
| ACTG1 | 1.26 | 0.015 |
| PALLD | 1.26 | 0.028 |
| CD63 | 1.24 | 0.005 |
| AKAP13 | 1.24 | 0.037 |
| GLUL | 1.24 | 0.018 |
| TRIM8 | 1.24 | 0.022 |
| JUN | 1.24 | 0.014 |
| BMPR2 | 1.24 | 0.040 |
| LGALS3 | 1.24 | 0.013 |
| CRIP2 | 1.24 | 0.039 |
| ANXA1 | 1.23 | 0.010 |
| NPC2 | 1.22 | 0.023 |
| CRTAP | 1.22 | 0.037 |

**Table S2. Upregulated genes on the migration trails in the ovarian cancer sample**

|  | **Fold Change** | **Adjusted P values** |
| --- | --- | --- |
| IL1RN | 2.29 | 0.039 |
| PLA2G5 | 2.26 | 0.083 |
| ITM2A | 2.13 | 0.016 |
| PDLIM3 | 2.13 | 0.021 |
| ADIRF | 2.11 | 0.032 |
| COL23A1 | 2.10 | 0.000 |
| ADM | 2.00 | 0.010 |
| CSTA | 1.94 | 0.000 |
| FHOD3 | 1.90 | 0.088 |
| MYCN | 1.89 | 0.059 |
| HSPB6 | 1.85 | 0.088 |
| SIM2 | 1.84 | 0.086 |
| ZNF469 | 1.84 | 0.078 |
| NDUFA4L2 | 1.83 | 0.000 |
| RAMP3 | 1.81 | 0.083 |
| BICC1 | 1.81 | 0.060 |
| STMN3 | 1.80 | 0.029 |
| MEX3B | 1.77 | 0.042 |
| SOBP | 1.76 | 0.052 |
| SLC11A1 | 1.75 | 0.000 |
| HNMT | 1.73 | 0.010 |
| ANK2 | 1.72 | 0.069 |
| PCED1B | 1.72 | 0.039 |
| COL5A1 | 1.72 | 0.069 |
| MRC1 | 1.72 | 0.042 |
| NEURL1B | 1.71 | 0.000 |
| SYTL2 | 1.69 | 0.035 |
| SCN7A | 1.67 | 0.052 |
| MPPED2 | 1.66 | 0.024 |
| GADD45B | 1.64 | 0.090 |
| MTSS1 | 1.62 | 0.000 |
| MN1 | 1.61 | 0.066 |
| ABCA6 | 1.61 | 0.060 |
| MBNL3 | 1.59 | 0.000 |
| HSPA1A | 1.58 | 0.000 |
| VCAN | 1.58 | 0.039 |
| CHL1 | 1.56 | 0.071 |
| RSAD2 | 1.56 | 0.080 |
| BTBD19 | 1.55 | 0.073 |
| CCN2 | 1.55 | 0.071 |
| SYNPO2 | 1.54 | 0.092 |
| SYNPO | 1.54 | 0.058 |
| GLT8D2 | 1.53 | 0.039 |
| COL5A2 | 1.53 | 0.080 |
| CCDC80 | 1.52 | 0.080 |
| TNXB | 1.52 | 0.088 |
| CNRIP1 | 1.51 | 0.065 |
| MPEG1 | 1.51 | 0.016 |
| G0S2 | 1.51 | 0.000 |
| TMEM131L | 1.51 | 0.010 |
| CCL3 | 1.50 | 0.070 |
| HSH2D | 1.50 | 0.042 |
| GDF15 | 1.49 | 0.032 |
| NUPR1 | 1.49 | 0.049 |
| HSPA6 | 1.49 | 0.059 |
| MIR1915HG | 1.48 | 0.074 |
| NCF2 | 1.48 | 0.029 |
| WNT5A | 1.48 | 0.049 |
| PLEK | 1.47 | 0.058 |
| GRIK2 | 1.47 | 0.024 |
| CELF2 | 1.46 | 0.074 |
| PDLIM5 | 1.46 | 0.010 |
| PPP1R15A | 1.46 | 0.010 |
| SECTM1 | 1.45 | 0.095 |
| LIFR | 1.45 | 0.010 |
| SPP1 | 1.45 | 0.064 |
| ALOX5AP | 1.45 | 0.010 |
| IFFO1 | 1.44 | 0.100 |
| KANK1 | 1.44 | 0.042 |
| TBX3 | 1.44 | 0.053 |
| GPR183 | 1.43 | 0.073 |
| ZEB1 | 1.43 | 0.085 |
| ZSWIM5 | 1.42 | 0.043 |
| GPR157 | 1.42 | 0.033 |
| TMEM132C | 1.42 | 0.067 |
| BHLHE40 | 1.42 | 0.010 |
| FRMD6 | 1.42 | 0.033 |
| LGALS3 | 1.41 | 0.010 |
| PPP1R14A | 1.41 | 0.056 |
| TRIB3 | 1.40 | 0.016 |
| HERC6 | 1.40 | 0.028 |
| VEGFA | 1.40 | 0.029 |
| PFKFB4 | 1.39 | 0.074 |
| F3 | 1.39 | 0.060 |
| SIGLEC1 | 1.39 | 0.016 |
| ANPEP | 1.39 | 0.085 |
| EGLN3 | 1.39 | 0.061 |
| AGFG2 | 1.38 | 0.024 |
| SSBP2 | 1.38 | 0.039 |
| SGCE | 1.37 | 0.050 |
| SPON2 | 1.37 | 0.000 |
| ASCL2 | 1.37 | 0.016 |
| FN1 | 1.36 | 0.094 |
| PIM1 | 1.36 | 0.024 |
| ITGA6 | 1.36 | 0.068 |
| RBPMS | 1.36 | 0.083 |
| PPFIA4 | 1.36 | 0.082 |
| GPNMB | 1.36 | 0.057 |
| SNCA | 1.35 | 0.042 |
| PPM1K | 1.35 | 0.035 |
| S100A9 | 1.35 | 0.029 |
| SLC2A1 | 1.35 | 0.066 |
| TIMP2 | 1.35 | 0.060 |
| DTNA | 1.34 | 0.000 |
| CLIC6 | 1.34 | 0.029 |
| PXDN | 1.34 | 0.010 |
| SAMD9 | 1.34 | 0.032 |
| RND3 | 1.33 | 0.000 |
| NMRK1 | 1.33 | 0.058 |
| EPB41L3 | 1.33 | 0.050 |
| NDRG1 | 1.32 | 0.000 |
| MAPRE2 | 1.32 | 0.039 |
| PLK2 | 1.32 | 0.074 |
| PFKFB3 | 1.32 | 0.028 |
| TNFSF10 | 1.32 | 0.039 |
| SPI1 | 1.31 | 0.032 |
| CCNA1 | 1.31 | 0.028 |
| CIART | 1.31 | 0.054 |
| HERC5 | 1.31 | 0.063 |
| PLSCR4 | 1.31 | 0.088 |
| FCGR2A | 1.31 | 0.061 |
| AMT | 1.31 | 0.091 |
| IFI44 | 1.30 | 0.069 |
| RAI14 | 1.30 | 0.028 |
| PRKD1 | 1.30 | 0.065 |
| APOL1 | 1.30 | 0.041 |
| KCNQ3 | 1.30 | 0.073 |
| PKD2 | 1.30 | 0.016 |
| KRT5 | 1.29 | 0.016 |
| CHSY1 | 1.29 | 0.000 |
| DIRAS3 | 1.29 | 0.053 |
| PALLD | 1.29 | 0.071 |
| COTL1 | 1.29 | 0.028 |
| TRIM21 | 1.29 | 0.070 |
| RCAN2 | 1.29 | 0.091 |
| PTPRG | 1.28 | 0.000 |
| CBX7 | 1.28 | 0.072 |
| DHX58 | 1.28 | 0.057 |
| SP110 | 1.28 | 0.068 |
| VIM | 1.28 | 0.077 |
| PLSCR1 | 1.27 | 0.055 |
| PHACTR2 | 1.27 | 0.033 |
| BOC | 1.27 | 0.052 |
| FXYD1 | 1.27 | 0.068 |
| SEMA3B | 1.27 | 0.060 |
| FAM110C | 1.27 | 0.032 |
| MOB3A | 1.27 | 0.000 |
| PIM3 | 1.26 | 0.010 |
| HK2 | 1.26 | 0.090 |
| ZHX3 | 1.26 | 0.024 |
| ATF7 | 1.26 | 0.047 |
| PLPPR2 | 1.25 | 0.054 |
| SUSD6 | 1.25 | 0.000 |
| ARHGDIB | 1.24 | 0.045 |
| ARMCX1 | 1.24 | 0.093 |
| FSTL3 | 1.24 | 0.053 |
| STAT6 | 1.24 | 0.042 |
| ARHGAP10 | 1.24 | 0.056 |
| RAB31 | 1.24 | 0.100 |
| MAGEH1 | 1.24 | 0.043 |
| AMPD3 | 1.24 | 0.060 |
| ZNF703 | 1.24 | 0.028 |
| ADAM8 | 1.24 | 0.093 |
| DHRS7 | 1.23 | 0.077 |
| PALM2-AKAP2 | 1.23 | 0.065 |
| ANKRD9 | 1.23 | 0.033 |
| COL27A1 | 1.23 | 0.038 |
| CD4 | 1.23 | 0.088 |
| NIBAN2 | 1.23 | 0.010 |
| C1orf21 | 1.23 | 0.088 |
| PSMB10 | 1.23 | 0.050 |
| TNFAIP2 | 1.22 | 0.088 |
| MTSS2 | 1.22 | 0.041 |
| NFIL3 | 1.22 | 0.045 |
| ASPH | 1.22 | 0.035 |
| JUNB | 1.22 | 0.068 |
| FERMT2 | 1.21 | 0.080 |
| AK1 | 1.21 | 0.059 |
| ATP6V0C | 1.21 | 0.000 |
| NEDD9 | 1.21 | 0.067 |
| SNX33 | 1.21 | 0.068 |
| MPDZ | 1.21 | 0.082 |
| PML | 1.21 | 0.055 |
| CFL2 | 1.21 | 0.067 |
| ENO2 | 1.20 | 0.060 |
| FTL | 1.20 | 0.000 |
| DNAJB1 | 1.20 | 0.000 |
| RXRA | 1.20 | 0.000 |
| ENGASE | 1.20 | 0.098 |
| INHBB | 1.20 | 0.033 |
| CNTFR | 1.20 | 0.068 |

**Table S3. Upregulated genes on the migration trails in the melanoma sample**

|  | **Fold change** | **Adjuste p values** |
| --- | --- | --- |
| C10orf99 | 6.65 | 0.000 |
| KRT6C | 5.82 | 0.000 |
| KRT14 | 4.14 | 0.000 |
| KRT75 | 3.95 | 0.000 |
| KRT5 | 3.35 | 0.000 |
| TYRP1 | 3.30 | 0.031 |
| MMP1 | 3.24 | 0.036 |
| GJB6 | 3.07 | 0.017 |
| KRT16 | 2.96 | 0.000 |
| EPGN | 2.88 | 0.017 |
| SFN | 2.85 | 0.010 |
| FGFBP1 | 2.75 | 0.026 |
| KRT6B | 2.69 | 0.010 |
| COL17A1 | 2.63 | 0.026 |
| MGST1 | 2.60 | 0.010 |
| FAM83C | 2.57 | 0.026 |
| IVL | 2.49 | 0.013 |
| SERPINB5 | 2.46 | 0.025 |
| PKP3 | 2.45 | 0.000 |
| MARCO | 2.45 | 0.048 |
| KRT17 | 2.27 | 0.034 |
| PKP1 | 2.25 | 0.019 |
| LAD1 | 2.24 | 0.041 |
| SOX15 | 2.20 | 0.034 |
| FRRS1 | 2.05 | 0.022 |
| JUP | 2.02 | 0.031 |
| SDC1 | 1.80 | 0.031 |
| USP18 | 1.78 | 0.000 |
| SPINT2 | 1.76 | 0.049 |
| CLEC11A | 1.76 | 0.036 |
| MX1 | 1.75 | 0.028 |
| PEX3 | 1.75 | 0.030 |
| DAAM2 | 1.73 | 0.030 |
| MAOB | 1.68 | 0.041 |
| S100A10 | 1.67 | 0.047 |
| IFIT1 | 1.65 | 0.049 |
| HERC5 | 1.65 | 0.041 |
| MSRB3 | 1.62 | 0.000 |
| PARP12 | 1.56 | 0.028 |
| DDX58 | 1.55 | 0.022 |
| MYD88 | 1.54 | 0.000 |
| OR51B5 | 1.54 | 0.000 |
| RAPGEF2 | 1.51 | 0.043 |
| CEACAM1 | 1.51 | 0.042 |
| CSTB | 1.49 | 0.013 |
| SLC22A23 | 1.45 | 0.006 |
| ENTHD1 | 1.45 | 0.000 |
| CDH12 | 1.45 | 0.000 |
| KIAA1522 | 1.41 | 0.013 |
| PAG1 | 1.41 | 0.019 |
| PRSS51 | 1.40 | 0.000 |
| TUBB2A | 1.36 | 0.042 |
| EIF2AK2 | 1.36 | 0.010 |
| MELTF | 1.35 | 0.019 |
| SH3BP5 | 1.35 | 0.000 |
| CBLB | 1.34 | 0.000 |
| DTX3L | 1.34 | 0.033 |
| IL13RA1 | 1.34 | 0.045 |
| CEP170 | 1.34 | 0.041 |
| MT1X | 1.30 | 0.019 |
| PSPH | 1.30 | 0.000 |
| MAMDC2 | 1.29 | 0.006 |
| BOP1 | 1.27 | 0.000 |
| PFN2 | 1.27 | 0.000 |
| TRIM63 | 1.27 | 0.000 |
| FERMT2 | 1.27 | 0.000 |
| MAPK6 | 1.27 | 0.026 |
| ACTN1 | 1.26 | 0.026 |
| RASAL2 | 1.26 | 0.000 |
| HK1 | 1.26 | 0.043 |
| LIMA1 | 1.26 | 0.044 |
| ALDH1B1 | 1.25 | 0.006 |
| PARP10 | 1.22 | 0.039 |
| SBNO2 | 1.22 | 0.037 |
| NUCB2 | 1.21 | 0.045 |
| KDELR3 | 1.21 | 0.010 |
| ADGRG6 | 1.21 | 0.033 |
| LGALS3BP | 1.19 | 0.030 |
| LY6K | 1.19 | 0.026 |
| B4GALT5 | 1.18 | 0.026 |
| H3F3B | 1.17 | 0.042 |
| BRMS1 | 1.17 | 0.000 |
| ARL2 | 1.17 | 0.000 |
| PRMT3 | 1.17 | 0.019 |
| GSTP1 | 1.16 | 0.000 |
| CD99 | 1.16 | 0.006 |
| AKAP12 | 1.15 | 0.017 |
| MT-ND6 | 1.14 | 0.045 |
| PARVA | 1.14 | 0.010 |
| EGLN3 | 1.14 | 0.026 |
| RRS1 | 1.14 | 0.006 |
| S100B | 1.14 | 0.010 |
| BCL2 | 1.14 | 0.043 |
| VARS | 1.14 | 0.000 |
| SGCD | 1.13 | 0.049 |
| PACS1 | 1.13 | 0.013 |
| RCN1 | 1.13 | 0.033 |
| HPS4 | 1.13 | 0.037 |
| FABP5 | 1.12 | 0.000 |
| STXBP6 | 1.11 | 0.041 |
| YAP1 | 1.10 | 0.019 |
